# Evaluating speech-like distortion product otoacoustic emissions: a comparison of time- and frequency-domain metrics across three datasets

**DOI:** 10.64898/2026.11.11.765899

**Authors:** Janna Steinebach, Tobias Reichenbach

**Affiliations:** Department Artificial Intelligence in Biomedical Engineering, Friedrich-Alexander-Universität Erlangen-Nürnberg, Nürnberger Strasse 74, 91052 Erlangen, Germany

**Keywords:** otoacoustic emissions, distortion product otoacoustic emissions, speech, cochlear mechanics, cross-correlation, magnitude-squared coherence, stimulus design

## Abstract

Speech-like distortion-product otoacoustic emissions (DPOAEs) allow cochlear activity to be measured during the perception of natural, continuous speech and have been used to study attentional effects in the cochlea. The technique is still young, however, and existing studies differ in stimulus design and in the metric used to quantify the emissions, which makes their results difficult to compare. Here, we reanalysed three existing datasets (106 participants, 14 stimulus types) with a common pipeline. We compared six metrics: the band power of the recording, the magnitude-squared coherence (MSC) between the recording and a DPOAE template, and four metrics derived from their complex cross-correlation, namely peak magnitude, signal-to-noise ratio (SNR), peak area and peak width. A separability index quantified how well each metric separated in-ear recordings from out-of-ear control recordings without emissions. Band power, MSC and SNR separated best, with mean separability indices of about 0.9, whereas the peak magnitude performed worst and was inverted for half of the stimulus types. Since band power does not require a DPOAE template, it serves as a sanity check of the setup rather than as evidence for an emission. The MSC was insensitive to the latency of the emission but well suited for comparisons across stimuli, whereas the cross-correlation additionally provided the latency. The emissions differed significantly between the stimulus types of the three datasets, and those below about 700 Hz were measured least reliably. We derive practical recommendations for the analysis and stimulus design of future experiments.

## 1 INTRODUCTION

The cochlea is a fascinating organ that does not merely detect sound, but actively amplifies it. Outer hair cells provide a frequency-specific mechanical gain, and as a by-product of the underlying nonlinearity, the cochlea emits sound that can be recorded with a sensitive microphone in the ear canal [1]. These otoacoustic emissions (OAEs) offer a non-invasive window onto cochlear function. Distortion-product OAEs (DPOAEs), evoked by two pure tones of frequencies *f*_1_ and *f*_2_ and measured at distortion frequencies such as 2*f*_1_ − *f*_2_, are routinely used in hearing research and in clinical practice, for instance in newborn hearing screening [2, 3]. Because the cochlear amplifier is controlled by the brain through the medial olivocochlear (MOC) efferent system [4], DPOAEs have also been used to investigate whether cognitive processes such as selective attention reach down to the cochlea. The results of these studies have, however, been inconsistent [5–8].

One reason for these inconsistencies may be the large variability in attentional tasks. Furthermore, the studies mentioned above used pure tone DPOAE measurements, and these are far from the sounds to which humans typically attend. To measure cochlear activity in response to natural speech, Saiz-Alía et al. [9] developed speech-like DPOAEs. Their stimuli are derived from the harmonic structure of voiced speech: they follow two harmonics of a talker’s time-varying fundamental frequency, and the resulting emission follows a third harmonic. Since the frequencies of stimuli and emission change continuously over time, speech-like DPOAEs cannot be read off a power spectrum like conventional DPOAEs. Instead, they were measured by cross-correlating the microphone recording with the expected waveform of the emission. Using this approach, speech-like DPOAEs have since been recorded in three datasets with different stimulus designs. Their attentional modulation, however, has been inconsistent as well: a small effect for one of two voices in the first study [9], robust effects for stimuli derived from resolved harmonics in a second study [10], and no consistent effects in a third [11].

Before such differences can be interpreted, it has to be established how reliably the analysis captures the emission in the first place, and how this depends on the stimuli. This question has not been addressed systematically so far. Speech-like DPOAEs are a young technique, and each study has relied on a single metric, the magnitude of the cross-correlation peak, either directly [10] or in the form of an attentional modulation coefficient [9]. Yet the recorded emissions differ considerably depending on stimulus properties, for instance in the width and the strength of their cross-correlation peaks, and a metric that works well for one design may not be suited for another. An established set of analysis methods, and guidance on when to use which, is still missing.

In this work, we present and compare different approaches to evaluate speech-like DPOAEs, in the time domain as well as in the frequency domain. In the time domain, we extend the cross-correlation analysis by several metrics that are derived from the cross-correlation peak: its magnitude, its signal-to-noise ratio, the area under the peak, and its width. In the frequency domain, we consider the band power of the recording and introduce the magnitude-squared coherence between the recording and the expected waveform of the emission. We apply all metrics to the three existing datasets, which together comprise 14 stimulus types and 106 participants, and we assess how reliably each metric separates in-ear recordings from out-of-ear control recordings. Based on these results, we derive recommendations both for the choice of metric and for the design of stimuli. Since we hope to make speech-like DPOAEs accessible to researchers who may not be familiar with the underlying signal processing, we explain all methods from their basic principles, and make the code openly available.

## 2 MATERIALS AND METHODS

### 2.1 Datasets

All analyses in this work are applied to three datasets of speech-like DPOAEs that were collected in our group since 2021 [9–11]. We refer to them as MSA21, JS25 and JS26. All three follow the same basic paradigm: participants listen to two competing talkers, one female and one male, that are presented to one ear, while speech-like DPOAEs are elicited and recorded in the contralateral ear (Fig. 1). Participants attended either the female or the male voice. All three datasets also generate their stimuli in the same way. They differ, however, in the voices, in the harmonics from which the stimuli were derived, in the number of DPOAE stimuli presented at the same time, and in how the attentional conditions were arranged. An overview is given in Table 1, and the individual DPOAE stimulus types are listed in Table 2. Fig. 1 illustrates the conceptual setup that is common to all three datasets; details such as the ear to which speech and stimuli were presented or the number of stimulus pairs presented at the same time differ between the datasets and are described in Sections 2.1.2 to 2.1.4.

**Table 1.**
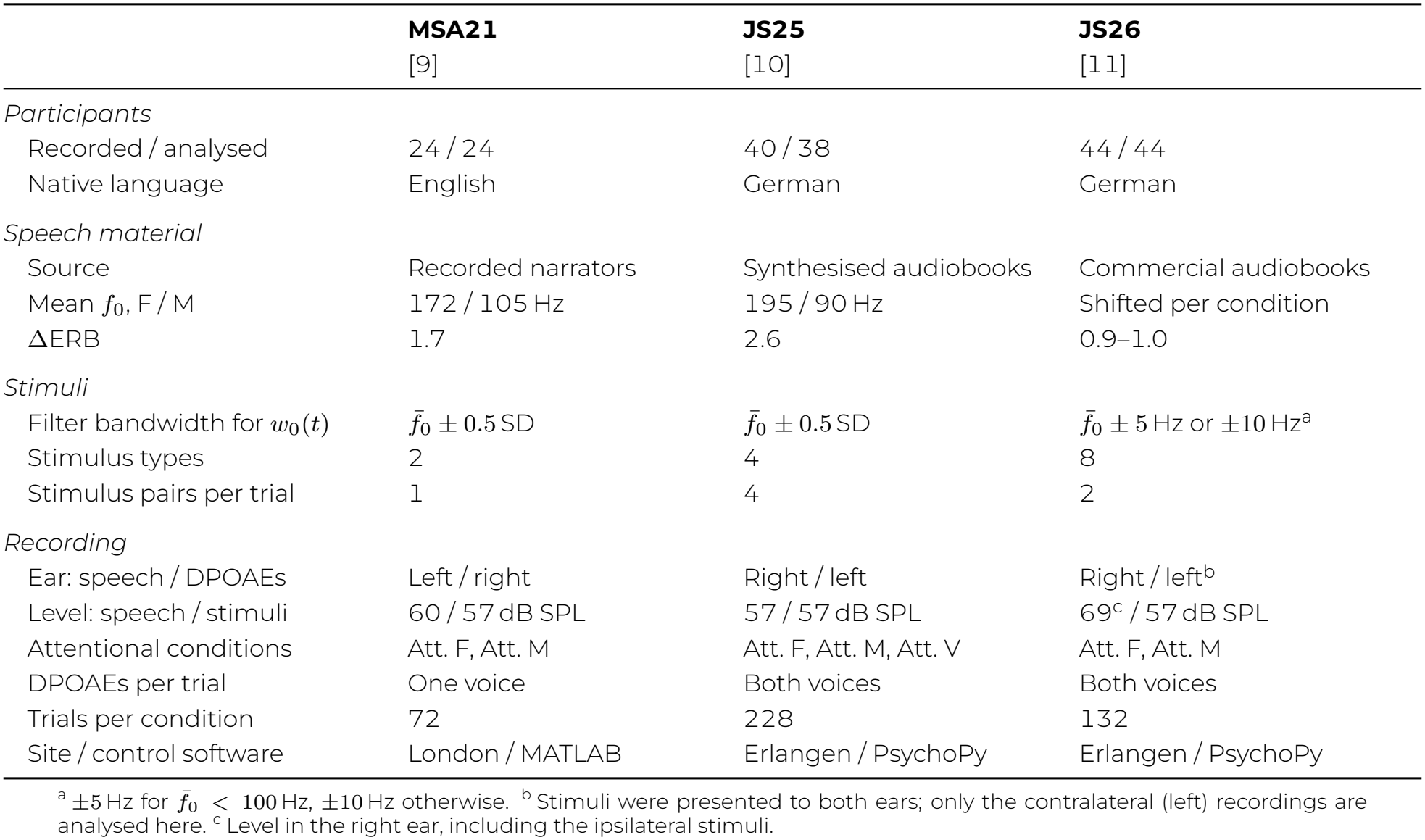
Overview of the three datasets. All trials lasted about two minutes. F: female voice; M: male voice; Att.: attended. The individual stimulus types of each dataset are listed in Table 2.

|  | **MSA21** | **JS25** | **JS26** |
|---|---|---|---|
|  | [9] | [10] | [11] |
| *Participants* |  |  |  |
| Recorded / analysed | 24 / 24 | 40 / 38 | 44 / 44 |
| Native language | English | German | German |
| *Speech material* |  |  |  |
| Source | Recorded narrators | Synthesised audiobooks | Commercial audiobooks |
| Mean *f*<sub>0</sub>, F / M | 172 / 105 Hz | 195 / 90 Hz | Shifted per condition |
| ΔERB | 1.7 | 2.6 | 0.9–1.0 |
| *Stimuli* |  |  |  |
| Filter bandwidth for *w*<sub>0</sub>(*t*) | *f̄*<sub>0</sub> ± 0.5 SD | *f̄*<sub>0</sub> ± 0.5 SD | *f̄*<sub>0</sub> ± 5 Hz or ±10 Hz<sup>a</sup> |
| Stimulus types | 2 | 4 | 8 |
| Stimulus pairs per trial | 1 | 4 | 2 |
| *Recording* |  |  |  |
| Ear: speech / DPOAEs | Left / right | Right / left | Right / left<sup>b</sup> |
| Level: speech / stimuli | 60 / 57 dB SPL | 57 / 57 dB SPL | 69<sup>c</sup> / 57 dB SPL |
| Attentional conditions | Att. F, Att. M | Att. F, Att. M, Att. V | Att. F, Att. M |
| DPOAEs per trial | One voice | Both voices | Both voices |
| Trials per condition | 72 | 228 | 132 |
| Site / control software | London / MATLAB | Erlangen / PsychoPy | Erlangen / PsychoPy |

<sup>a</sup> ±5 Hz for *f̄*<sub>0</sub> < 100 Hz, ±10 Hz otherwise. <sup>b</sup> Stimuli were presented to both ears; only the contralateral (left) recordings are analysed here. <sup>c</sup> Level in the right ear, including the ipsilateral stimuli.

**Table 2.**
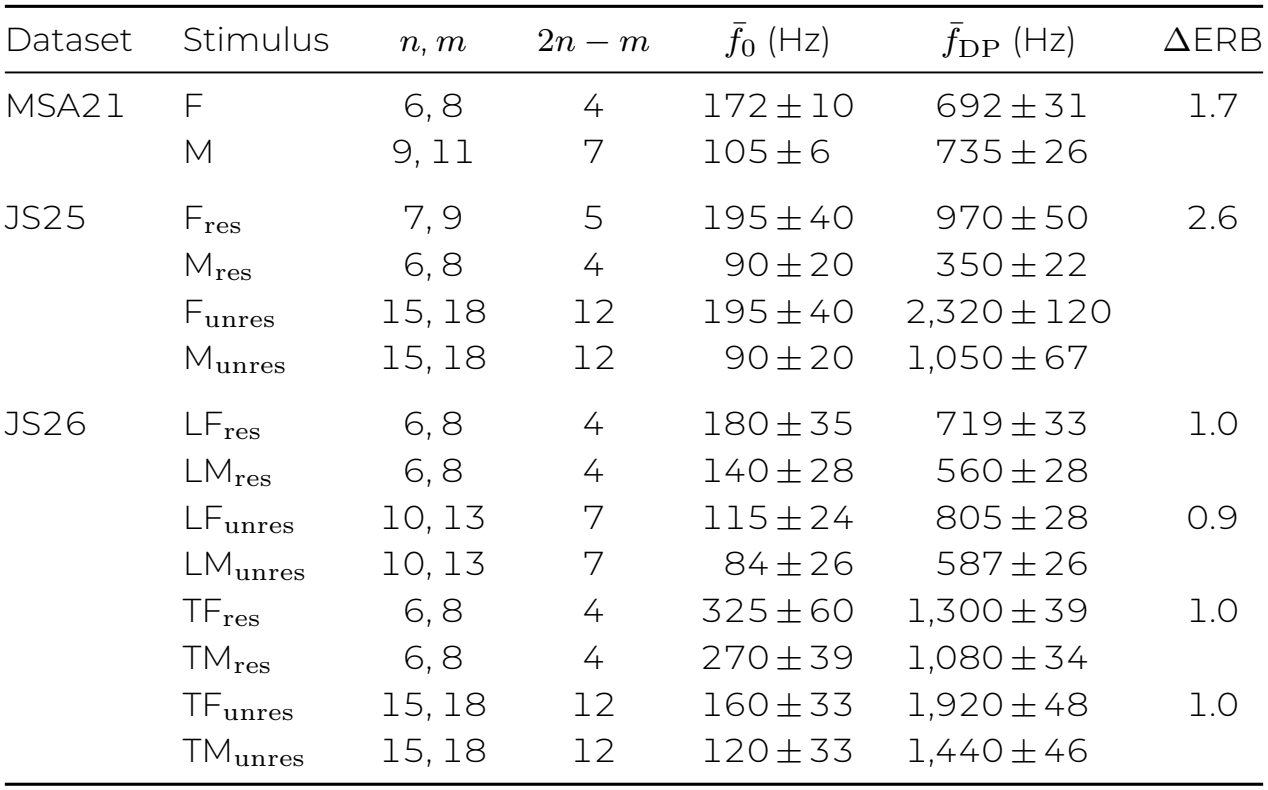
The 14 stimulus types of the three datasets, with the labels used in the figures. *n, m*: harmonic numbers of the stimulus pair; 2*n −m*: harmonic number of the DPOAE template; 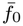: mean fundamental frequency of the voice (*±*SD across all chapters); 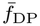: mean frequency of the DPOAE template ( *±*SD across all chapters). The SD of 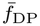 is considerably smaller than (2*n− m*) times the SD of 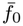, since the bandwidth *B*_0_ of the fundamental waveform (Table 1) limits how closely stimuli and templates follow the pitch contour. In JS26, the fundamental frequencies were shifted separately for each condition. L/T: low/transitional frequency regime; res/unres: derived from resolved/unresolved harmonics.

| Dataset | Stimulus | *n*, *m* | 2*n* − *m* | $\bar{f}_0$ (Hz) | $\bar{f}_{\mathrm{DP}}$ (Hz) | ΔERB |
|---|---|---|---|---|---|---|
| MSA21 | F | 6, 8 | 4 | 172 ± 10 | 692 ± 31 | 1.7 |
|  | M | 9, 11 | 7 | 105 ± 6 | 735 ± 26 |  |
| JS25 | F<sub>res</sub> | 7, 9 | 5 | 195 ± 40 | 970 ± 50 | 2.6 |
|  | M<sub>res</sub> | 6, 8 | 4 | 90 ± 20 | 350 ± 22 |  |
|  | F<sub>unres</sub> | 15, 18 | 12 | 195 ± 40 | 2,320 ± 120 |  |
|  | M<sub>unres</sub> | 15, 18 | 12 | 90 ± 20 | 1,050 ± 67 |  |
| JS26 | LF<sub>res</sub> | 6, 8 | 4 | 180 ± 35 | 719 ± 33 | 1.0 |
|  | LM<sub>res</sub> | 6, 8 | 4 | 140 ± 28 | 560 ± 28 |  |
|  | LF<sub>unres</sub> | 10, 13 | 7 | 115 ± 24 | 805 ± 28 | 0.9 |
|  | LM<sub>unres</sub> | 10, 13 | 7 | 84 ± 26 | 587 ± 26 |  |
|  | TF<sub>res</sub> | 6, 8 | 4 | 325 ± 60 | 1,300 ± 39 | 1.0 |
|  | TM<sub>res</sub> | 6, 8 | 4 | 270 ± 39 | 1,080 ± 34 |  |
|  | TF<sub>unres</sub> | 15, 18 | 12 | 160 ± 33 | 1,920 ± 48 | 1.0 |
|  | TM<sub>unres</sub> | 15, 18 | 12 | 120 ± 33 | 1,440 ± 46 |  |

**Figure 1:**
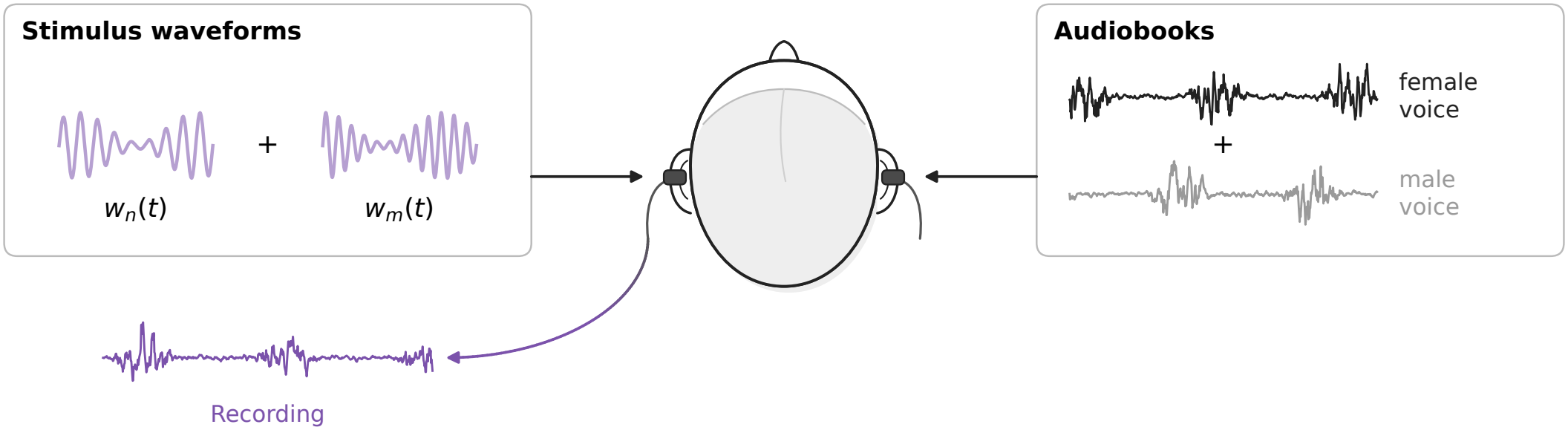
Conceptual illustration of the experimental setup. Participants were presented with two competing audiobooks, narrated by a female and a male voice, to one ear. An ear probe in the other ear delivered the stimulus waveforms *w*_*n*_(*t*) and *w*_*m*_(*t*) and recorded the sound pressure in the ear canal, which contains the speech-like DPOAE. The illustration is schematic; the ear of presentation and the number of simultaneously presented stimulus pairs differ between the datasets (Table 1).

We first summarise the DPOAE stimulus design shared by all three datasets. We recap it here, rather than only referring to the original publications, because the choices made at this stage determine what the recorded emissions look like and therefore which analysis is best suited to evaluate them (Sections 2.4 and 2.5).

#### 2.1.1 DPOAE stimulus design

Most parts of speech are voiced, that is, they are produced by vibration of the vocal folds. Voiced speech has a harmonic structure: its energy is concentrated at a fundamental frequency *f*_0_(*t*) and at the integer multiples *nf*_0_(*t*) of it, the harmonic overtones. In contrast to the fixed primary frequencies of a conventional DPOAE measurement, *f*_0_(*t*) changes continuously over time, typically within a range of some tens of hertz around the talker’s mean fundamental frequency 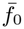.

Speech-like DPOAEs make use of this structure [9]. For each voiced segment of a speech signal, we first computed a fundamental waveform *w*_0_(*t*) that oscillates at *f*_0_(*t*). It was obtained by bandpass filtering the speech signal around 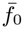 with a zero-phase Butterworth filter, and was normalised by z-scoring. The bandwidth *B*_0_ of this filter is a design parameter to which we return below. Using the Hilbert transform ℋ, we then computed the analytic representation of the fundamental waveform,

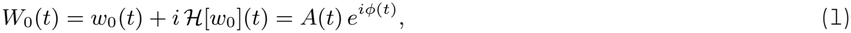

with the instantaneous amplitude *A*(*t*) = |*W*_0_(*t*)| and the instantaneous phase *ϕ*(*t*) = arg *W*_0_(*t*). A waveform that follows the *n*-th harmonic was then obtained by multiplying the phase by *n* and taking the real part of the signal [9, 10]:

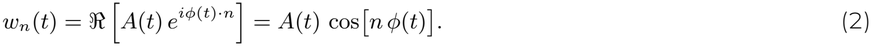

Its instantaneous frequency is *nf*_0_(*t*), and its amplitude follows the voicing of the speech signal: it is zero during un-voiced parts and pauses.

Two such waveforms *w*_*n*_(*t*) and *w*_*m*_(*t*) with *n < m* form a stimulus pair. They take the role of the primary tones *f*_1_ and *f*_2_ of a conventional DPOAE measurement and were delivered to the ear canal through two separate loudspeakers of the probe to avoid distortion in the hardware. Their lower-sideband cubic distortion product emerges at the instantaneous frequency (2*n* − *m*)*f*_0_(*t*), which corresponds to the waveform *w*_2*n−m*_(*t*), computed with Eq. (2) as well. We refer to *w*_2*n−m*_(*t*) as the template of the speech-like DPOAE. Because the template is known exactly, the recording can be compared against it both in the time domain (Section 2.4) and in the frequency domain (Section 2.5).

The bandwidth *B*_0_ of the filter that extracts *w*_0_(*t*) affects the resulting speech-like DPOAEs. Since *w*_*n*_(*t*) is obtained by multiplying the phase of *w*_0_(*t*) by *n*, the frequency fluctuations of *w*_*n*_(*t*), and therefore its bandwidth, scale approximately with *nB*_0_. A wide filter preserves more of the natural variation of *f*_0_(*t*), which results in broadband stimuli and templates. A narrow filter restricts *f*_0_(*t*) to a small range around 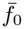, which results in more clearly determined stimulus frequencies, but comes with a loss of natural variability. As we show in Section 2.4.2, this directly determines the temporal resolution that can be achieved with cross-correlation.

Following previous work [9, 10], we distinguish between stimuli that were derived from resolved and from unresolved harmonics. Harmonics up to about the 10th are generally considered to be resolved by the cochlea, that is, they cause separate peaks along the basilar membrane, while higher harmonics overlap [12, 13]. As in [10], the terms refer to the harmonic structure of the speech signal from which a stimulus pair was derived, not to the two waveforms of the pair themselves, which are separated by a similar frequency ratio in all cases.

To describe how far apart the two competing voices are, we computed the difference of their mean fundamental frequencies on the ERB-number scale [14],

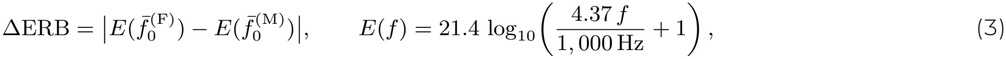

in which 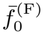 and 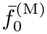denote the mean fundamental frequencies of the female and the male voice. ΔERB measures the distance between the two voices in units of auditory filter bandwidths, that is, in a way that reflects how far apart their fundamental frequencies are represented in the cochlea. We report it here as a descriptive feature of the stimuli and return to it in the Discussion.

#### 2.1.2 Dataset MSA21

Dataset MSA21 was collected by Saiz-Alía and colleagues and is described in detail in [9]. It comprises recordings from 24 native English speakers (14 female, 10 male, aged 18 to 26 years). The speech material consisted of two stories read by a male and by a female narrator, respectively, with mean fundamental frequencies of 105 ± 6 Hz and 172 ± 10 Hz (mean ± SD), which corresponds to ΔERB = 1.7. The fundamental frequency was estimated with PRAAT [15], and the fundamental waveform was extracted with a sixth-order zero-phase filter with passband edges 0.5 standard deviations below and above 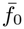.

Only one stimulus pair was presented at a time. For the male voice, the pair consisted of the 9th and 11th harmonics, yielding the template *w*_7_(*t*) at 735 ± 26 Hz. For the female voice, it consisted of the 6th and 8th harmonics, yielding *w*_4_(*t*) at 692 ± 31 Hz. The competing talkers were presented to the left ear at 60 dB SPL, and the stimulus waveforms were presented to the right ear at 57 dB SPL, where the emissions were recorded as well. The data were recorded at Imperial College London, with stimulus presentation and recording controlled by MATLAB.

In each two-minute trial of the competing-speaker task, speech-like DPOAEs were measured in relation to only one of the two voices, and participants attended either this voice or the competing one. Attended and ignored states were therefore measured in separate presentations of the same speech segment. For each voice, three such pairs of trials were recorded, which amounts to 12 trials per participant. Saiz-Alía et al. reported a moderate attentional modulation for the female voice, with larger speech-like DPOAEs when the female voice was attended, and no effect for the male voice [9]. However, to obtain this result, they excluded participants with low comprehension scores, which reduced the number of participants to 20 for the female and 19 for the male voice. In the present analysis, we included all 24 participants, since the key for the correct answers to the comprehension questions was not available for the present analysis.

#### 2.1.3 Dataset JS25

Dataset JS25 is described in detail in [10]. It comprises recordings from 40 native German speakers (21 female, 19 male, aged 18 to 31 years). As in the original publication, two participants were excluded, one because of comprehension scores below chance and one because of a faulty recording, so that 38 participants entered the present analyses. Two German audiobooks were synthesised with a text-to-speech engine, with a female voice of 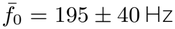 and a male voice of 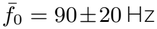. Their larger separation compared to MSA21 corresponds to ΔERB = 2.6. The fundamental frequency was estimated with the probabilistic YIN algorithm [16], and the filter for the fundamental waveform was the same as in MSA21.

The design was more ambitious than that of MSA21 in two respects. First, four stimulus pairs were presented simultaneously: one derived from resolved and one from unresolved harmonics of each voice (Table 2). This decision was based on the observation in MSA21 that attentional effects appeared only for the more resolved harmonics of the female voice.

All waveforms with the lower harmonic number were summed and played through one loudspeaker, and all waveforms with the higher harmonic number through the other. Second, the attentional state was varied while the acoustic input stayed the same: in every trial, both talkers and all four stimulus pairs were presented, and participants attended the female voice, the male voice, or a text on a screen. The attended state of one voice was therefore recorded at the same time as the ignored state of the other voice. Attended and ignored states of the same voice therefore were recorded with the same setup and the same acoustic input; only the attentional task changed. The speech was presented to the right ear and the stimuli were presented to and recorded from the left ear, both at an average of 57 dB SPL. These levels are 20 dB higher than those reported in the original publication [10]. The function that computed the average level of the recordings contained an error, so that the levels were underestimated there; it was corrected for the present work. The corrected levels are close to those of MSA21 and identical to those of JS26, which makes the three datasets more directly comparable. Stimulus presentation and recording were controlled by PsychoPy [17]. Each attentional condition was repeated 6 times, which amounts to 18 trials per participant.

Speech-like DPOAEs could be measured reliably for three of the four stimulus types using the cross-correlation approach, but not for M_res_, whose template had a mean frequency of only 350 Hz. Strong attentional effects emerged for F_res_, with smaller speech-like DPOAEs when the female voice was attended than when it was ignored. Effects also emerged for M_unres_, which were attributed to the frequency overlap of M_unres_ with F_res_ [10]. This overlap motivated the design of dataset JS26, which aimed at disentangling frequency region and resolvability.

#### 2.1.4 Dataset JS26

Dataset JS26 comprises recordings from 44 participants (22 female, 22 male, aged 20 to 43 years) and is described in detail in [11]. No participant had to be excluded. The study was designed to reproduce the attentional effects of JS25 and to investigate how they depend on the frequency region in which the stimuli are placed. Compared with JS25, the design was changed in four ways.

First, only two stimulus pairs were presented at the same time, one derived from each voice and both belonging to the same frequency and resolvability condition. This gave more freedom in choosing harmonics and fundamental frequencies without creating spectral overlap between simultaneously presented pairs.

Second, the stimulus pairs were constrained to presumed frequency regimes of cochlear mechanics: a low-frequency regime up to 1.5 kHz and a transitional regime between 1.5 and 4 kHz [18–22]. Crossing these with resolved and unresolved harmonics resulted in four conditions: low resolved, low unresolved, transitional resolved and transitional unresolved (Table 2).

Third, to place the harmonics of both voices in the intended regimes, the fundamental frequencies of the audiobooks were shifted separately for each condition. To this end, we used pitch-synchronous overlap-add (PSOLA) resynthesis in PRAAT [15], accessed through Parselmouth [23]. The *f*_0_(*t*) contour of each segment was shifted on a logarithmic scale such that its mean matched the target value, while its relative variability was preserved. Three iterations of resynthesis and re-estimation of the median *f*_0_ were used to correct for residual deviations from the target. The resulting mean fundamental frequencies are listed in Table 2. Because both voices had to fit into the same frequency regime, their fundamental frequencies were considerably closer than in the previous datasets, with ΔERB between 0.9 and 1.0. Unlike in JS25, the speech material consisted of commercial audiobooks read by professional narrators: “Den Hund überleben” [24], read by a male narrator, and “Eine Frau erlebt die Polarnacht” [25], read by a female narrator.

Fourth, the bandwidth *B*_0_ of the filter that extracts the fundamental waveform was restricted to a fixed, narrow band of 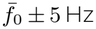 for 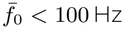 and 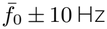 otherwise, instead of being set by the standard deviation of *f*_0_. The aim was to reduce correlations between the stimulus waveforms and the templates, which were stronger due to the closer fundamental frequencies, and thereby to improve the signal-to-noise ratio of the measurement. As we show in Section 2.4.2, this choice had a large effect on the temporal resolution of the cross-correlation.

As in JS25, all waveforms with the lower harmonic number were summed and presented through one loudspeaker, and those with the higher harmonic number through the other, and stimulus presentation and recording were controlled by PsychoPy [17]. Participants attended either the female or the male voice. Because speech-like DPOAEs related to both voices were elicited in every trial, the attended state of one voice was again recorded at the same time as the ignored state of the other voice, and the acoustic input was identical in both attentional conditions. Each trial lasted two minutes, and each combination of stimulus condition and attentional condition was presented three times, which resulted in 24 trials per participant. The audiobooks were presented to the right ear. The stimulus waveforms were presented to both ears, so that speech-like DPOAEs could be recorded simultaneously from the ear contralateral and from the ear ipsilateral to the speech. On the contralateral side, the average sound pressure level was 57 dB SPL, the same as in MSA21 and JS25. On the ipsilateral side, it was 69 dB SPL on average, mostly due to the louder audiobooks, which masked the presence of the ipsilateral stimulus waveforms. The present analyses use the contralateral recordings only, which are directly comparable to those of JS25. The evaluation of the ipsilateral recordings is still ongoing. Unlike in JS25, no consistent attentional effects were found in this dataset [11].

### 2.2 Common preprocessing

All recordings of the three datasets, including those that had been analysed before, were reanalysed with the same Python pipeline, so that all metrics are directly comparable across datasets. Recordings were sampled at 44.1 kHz and converted to sound pressure using the microphone sensitivity of 50 mV/Pa and the amplifier gain of 40 dB for JS25 and JS26 and of 0 dB for MSA21.

Before any further analysis, the delay introduced by the measurement equipment was removed from each trial. To this end, the recording was cross-correlated with each of the two stimulus waveforms, using the method described in Section 2.4.1. Since the stimulus waveforms are recorded directly by the microphone, these cross-correlations peak at the equipment delay. The two delays were averaged, and the recording was shifted by the result.

### 2.3 Out-of-ear recordings

For each dataset, 12 out-of-ear control recordings were obtained, mimicking 12 participants. The experimental procedure was exactly the same as for the respective participants of each dataset, except that the probe was not inserted into an ear canal. The probe could therefore not pick up any speech-like DPOAE. The recordings were processed with exactly the same pipeline as the in-ear recordings and serve as a measure of what each metric yields in the absence of an emission (Section 2.7).

### 2.4 Time-domain analysis via cross-correlation

Cross-correlation was established as the standard analysis of speech-like DPOAEs by Saiz-Alía et al. [9]. Its logic is simple. Conventional DPOAEs are read off a power spectrum at a fixed distortion frequency. For speech-like DPOAEs, this is not as straight-forward, because the distortion frequency (2*n* − *m*)*f*_0_(*t*) changes continuously over time. We are, however, in the fortunate position of knowing what the emission should look like: it should resemble the template *w*_2*n−m*_(*t*). The template can therefore be slid along the recording, and the similarity between the two can be computed for different possible delays. If the ear produces the emission with a certain delay, the similarity peaks at this delay. This procedure has a further benefit that none of the frequency-domain methods in Section 2.5 offer: it tells us how long it takes for the emission to appear in the ear canal.

#### 2.4.1 Complex cross-correlation

The emission can have an arbitrary phase shift with respect to the template. A plain cross-correlation would then vary with this phase and could even vanish. We therefore computed the complex cross-correlation between the recording *r*(*t*) and the analytic representation of the template [9]. In discrete time, with lag *τ*, its real and imaginary parts are

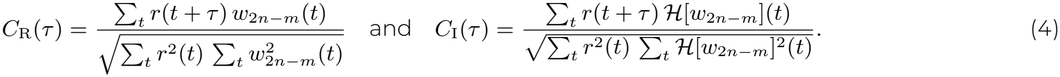

The normalisation ensures that the auto-correlation of a waveform equals one at zero lag, so that the values can be read as correlation coefficients. Positive lags correspond to the recording lagging behind the template. The envelope of the complex cross-correlation,

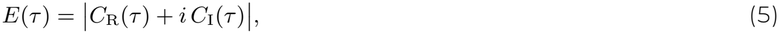

no longer depends on the phase of the emission. It was smoothed with a zero-phase third-order Butterworth low-pass filter, whose cut-off was set to the dominant frequency of *C*_R_(*τ* ) divided by a factor *c*. We used *c* = 10 for JS25 and JS26, and *c* = 2 and *c* = 0.1 for the female and the male voice of MSA21, respectively. In each case, the value was determined iteratively as the weakest smoothing that resulted in an envelope that no longer followed the oscillations of the carrier. Lags between −0.5 and 0.5 s were retained for further analysis.

The cross-correlation of the template with itself, its auto-correlation, shows what the envelope of a perfect, noise-free recording would look like (cf. Fig. 2A in [10]). Its width determines how sharply the emission can be located in time. We quantified it by the standard deviation *σ*_ac_ of a Gaussian with the same full width at half maximum as the central peak of the auto-correlation envelope, 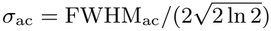.

**Figure 2:**
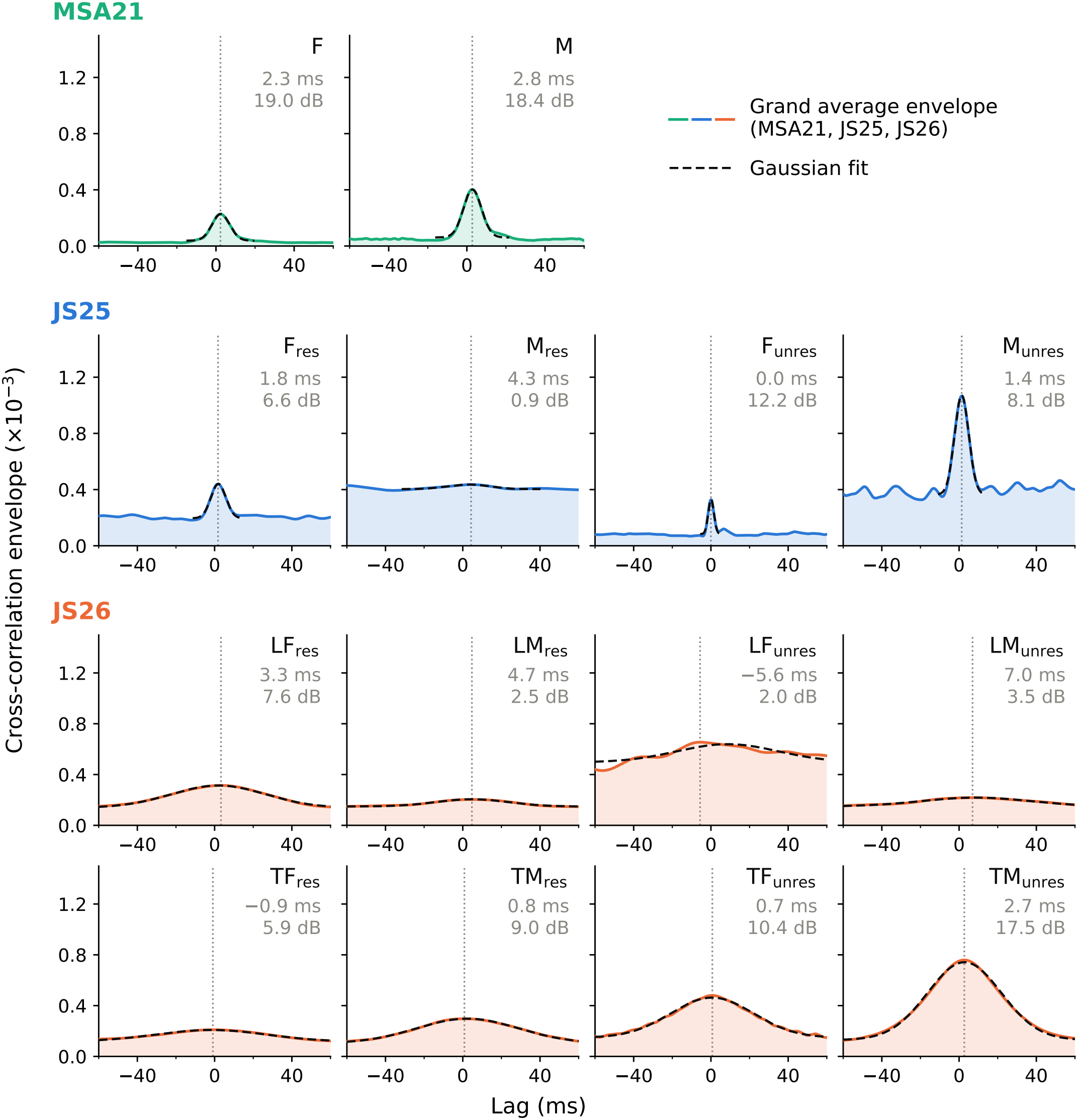
Grand average envelopes of the complex cross-correlation for all stimulus conditions of the three datasets, pooled across attentional conditions (coloured lines), together with the fitted Gaussians (Eq. 8, dashed black lines). The dotted vertical lines indicate the lag of the GA peak; the numbers in grey give this lag and the GA SNR. MSA21: female (F) and male (M) voice. JS25: stimuli derived from resolved (res) and unresolved (unres) harmonics of the female (F) and male (M) voice. JS26: as JS25, separately for the low (L) and transitional (T) frequency regime. The peaks of JS26 are an order of magnitude broader than those of MSA21 and JS25, reflecting the narrower bandwidth of the stimulus templates.

Speech-like DPOAEs are expected within a few milliseconds of zero lag [9, 10]. At much longer lags, the template and the recording are unrelated, and the envelope only reflects correlations that arise by chance. We therefore defined a noise region *N* of lags 100 ms ≤ |*τ* | ≤ 500 ms on both sides of the peak and used the mean envelope in this region as the noise floor:

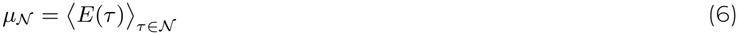

This noise floor is never zero. In the absence of an emission, *C*_R_ and *C*_I_ fluctuate randomly around zero. Their envelope *E*, however, is always positive and approximately follows a Rayleigh distribution, whose mean is proportional to the standard deviation of the fluctuations. Any measurement of the envelope therefore contains a positive offset that depends on the noise in the recording.

#### 2.4.2 Grand average cross-correlation

As a first step, we tested whether speech-like DPOAEs emerged consistently in a given condition. To this end, we averaged the envelopes *E*_*k*_(*τ* ) of all trials *k* = 1, …, *K* of all participants in this condition, which yields the grand average (GA)

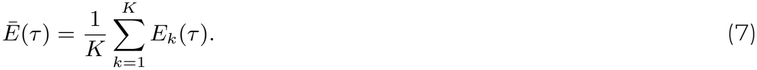

The GA peak was located as the maximum of *Ē*(*τ* ) within the latency range of ±15 ms. To describe its position and shape, we fitted a Gaussian with a constant baseline,

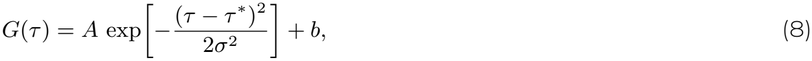

by nonlinear least squares within a window of ±4*σ*_ac_ around the peak, using *σ*_ac_ as starting value for *σ*. The fitted centre *τ*^*\**^ served as the latency of the speech-like DPOAE in this condition and as the reference lag for the metrics in Section 2.4.3. As a summary of the strength of the GA peak, we report the GA signal-to-noise ratio (SNR), computed as in Eq. (9) with the GA envelope *Ē*(*τ* ) in place of *E*_*k*_(*τ* ) and evaluated at the GA peak.

A GA peak was considered significant if it exceeded the maximum of *Ē*(*τ* ) in the noise region. This criterion can be understood as an empirical surrogate test. The noise region contains the cross-correlation of the recordings with the template shifted by 100 to 500 ms in either direction, that is, by lags at which no emission is expected. Each of these lags therefore provides a surrogate value of the envelope in the absence of an emission. If the noise region contained *N* independent surrogate values, the probability that the GA peak exceeds all of them by chance would be 1*/*(*N* + 1). Neighbouring lags, however, are not independent, since the envelope varies smoothly over a range set by the width of the auto-correlation of the template. The number of independent surrogate values is therefore approximately the total length of the noise region, 800 ms, divided by the width of the auto-correlation peak (about 4*σ*_ac_). For the narrow peaks of MSA21 and JS25, with *σ*_ac_ between 1.4 and 9 ms, this yields between about 20 and 140 independent values and false-positive rates between about 0.01 and 0.04. For the broad peaks of JS26, with *σ*_ac_ between 20 and 36 ms, it yields fewer than ten, which corresponds to a false-positive rate of about 0.1. The criterion therefore becomes lenient for broad, smooth envelopes. Indeed, all GA peaks of the present datasets passed it, including those that showed no clear peak (Fig. 2).

We therefore additionally required the GA peak to exceed the mean noise floor *μ*_*N*_ (Eq. 6) by at least four standard deviations of *Ē*(*τ* ) in the noise region. Unlike the maximum, the standard deviation does not depend on the number of independent values in the noise region. It also accounts for how strongly the GA fluctuates in a given condition, which a fixed threshold on the GA SNR would not: the relative standard deviation of the noise region ranged from 3 to 14 % across the conditions of the present datasets, so that the same GA SNR can stand for a clear peak in one condition and for a fluctuation of the noise in another. By Cantelli’s inequality [26], a value of any distribution exceeds its mean by four standard deviations with a probability of at most 1*/*(1 + 4^2^) ≈ 0.06, and for a Gaussian distribution with a probability of about 3 · 10^*−*5^. In the present data, the GA peaks lay either at most 3.6 or at least 6.5 standard deviations above the noise floor, so that the exact value of the threshold did not affect which conditions met the criterion.

Examples of the GAs for all three datasets are shown in Fig. 2. For this figure, the trials of the different attentional conditions were pooled. In all three datasets, the GA approach was valid and produced clear peaks for most conditions. The peaks were most pronounced in MSA21, with GA SNRs of 19.0 dB for the female and 18.4 dB for the male voice. In JS25, GA SNRs ranged from 6.6 dB (F_res_) to 12.2 dB (F_unres_), and in JS26 from 2.5 dB (LM_res_) to 17.5 dB (TM_unres_) for the conditions that met both criteria. Two conditions failed to produce a clear GA peak and did not meet the second criterion: M_res_ in JS25 (GA SNR of 0.9 dB, 3.6 standard deviations above the noise floor), as already reported in [10], and LF_unres_ in JS26 (2.0 dB, 2.6 standard deviations). All other GA peaks lay at least 6.5 standard deviations above the noise floor. For LF_unres_, the maximum of the GA envelope even lay at a negative lag of −5.6 ms, which would indicate that the emission preceded the stimulus. This is implausible and further limits the interpretability of this condition. In the present work, we nevertheless kept all conditions in the subsequent analyses, in order to show how the different metrics behave when no reliable emission can be measured.

The temporal resolution differed markedly between the datasets. In MSA21 and JS25, the GA peaks were narrow, with fitted widths of *σ* ≈ 4.5 ms in MSA21 and between 1.5 and 3.9 ms for the reliable conditions of JS25. In JS26, they were an order of magnitude broader, with *σ* between 19 and 29 ms. In all datasets, the fitted widths closely matched those of the auto-correlations of the templates, *σ*_ac_. The width of the peak is therefore set by the template rather than by temporal jitter of the emission, as already noted in [10]. In retrospect, we can attribute the poor temporal resolution of JS26 to the narrow bandwidth *B*_0_ that was chosen during stimulus design (Section 2.1.1). In MSA21 and JS25, *B*_0_ was set by the standard deviation of the talker’s *f*_0_, whereas for JS26, it was restricted to 10 or 20 Hz to reduce correlations between the stimulus waveforms and the templates. By the time–frequency uncertainty principle [27], a narrowband template necessarily has a long auto-correlation. A practical consequence concerns the noise region: with *σ* ≈ 25 ms, the tails of the peak extend to about 75 ms (3*σ*). We therefore let the noise region start at 100 ms for all datasets (Eq. 6), so that it does not contain parts of the peak, but still remains consistent across datasets.

#### 2.4.3 Metrics

The GA establishes that, and at which lag, speech-like DPOAEs occur in a given condition. To compare conditions, for instance attended and ignored states, a single metric that charaterizes the speech-like DPOAE for every participant would be valuable. We derived four such candidate metrics from the cross-correlation (Fig. 3). They are presented in the order in which they add information: the first two describe only the height of the envelope at the GA peak, the latter two also take its shape into account.

**Figure 3:**
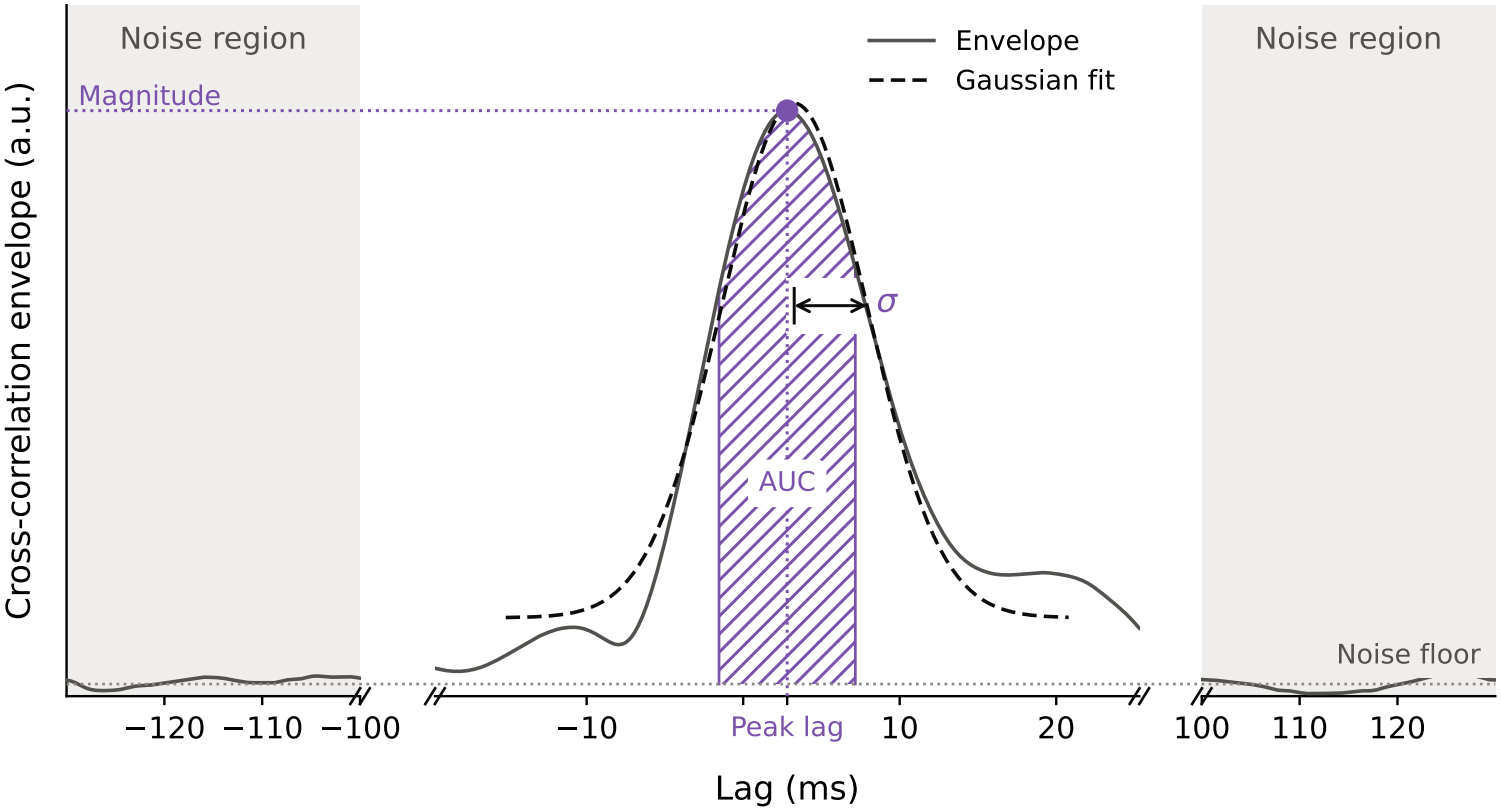
Metrics derived from the cross-correlation, illustrated on an exemplary speech-like DPOAE recording of the MSA21 dataset. The noise region (grey) at lags |*τ* |*≥* 100 ms defines the noise floor (dotted horizontal line). The peak magnitude is the value of the envelope at the peak lag, which in the analyses was taken from the grand average. The area under the curve (AUC, purple hatched) is computed above the noise floor within *±σ*_ac_ around the peak lag. The width *σ* is obtained from a Gaussian fit (dashed line). Note that the lag axis is interrupted on both sides of the peak.

All lag-dependent metrics were evaluated at the reference lag *τ*^*\**^ of the respective condition, obtained from the GA (Section 2.4.2). This avoids searching for a peak in every single trial, which would select the largest noise fluctuation whenever the emission is weak and thereby bias the values upwards. Metrics were reduced to one value per participant and condition in one of two ways. For metrics marked avgP, the metric was computed for every trial and then averaged across the trials of a participant. For metrics marked pP, the envelopes of all trials of a participant were first averaged into a participant-level GA, and the metric was computed from this average. The pP route is required for metrics that depend on the shape of the peak. Since this is quite variable, and can often not be inferred from a single trial due to excessive noise, a participant-level GA is used to provide robustness.

##### Peak magnitude (avgP)

The simplest metric is the value of the envelope at the reference lag, *E*_*k*_(*τ*^*\**^), averaged across the trials of a participant. This is the amplitude used in [10]. Its main limitation is the noise floor discussed in Section 2.4.1: since *E* is the magnitude of a complex number, the peak magnitude contains a positive contribution from noise even when no emission is present. Two recordings with the same emission but different noise levels therefore yield different magnitudes. As long as the noise is the same in the compared conditions, which is approximately the case when attended and ignored states are recorded with the same setup and the same acoustic input, this offset largely cancels in a comparison between the two attentional states. It does, however, compress relative differences between conditions, and it makes absolute values hard to compare across participants, sessions and datasets.

##### Signal-to-noise ratio (avgP)

To account for the noise floor, we computed for every trial the signal-to-noise ratio

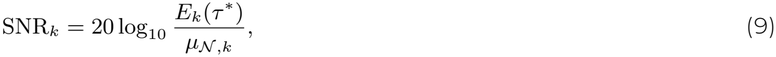

with the noise floor *μ*_*N,k*_ of the same trial (Eq. 6), and averaged it across the trials of a participant. Because each trial is referenced to its own noise, the SNR is more comparable across sessions and participants than the raw magnitude. It is, however, not necessarily superior in comparing attentional conditions. The noise floor does not only reflect acoustic noise in the ear canal: it also contains chance correlations between the template and everything else in the recording, in particular the stimulus waveforms and, in JS25 and JS26, the stimuli related to the competing voice. These correlations differ between templates, and thus between the DPOAEs related to the two voices. A difference in SNR can therefore arise from a difference in the noise floor rather than in the emission. When using the SNR as a metric, one therefore needs to consider what the noise region represents within the specific experimental design.

##### Area under the curve (pP)

For narrow peaks, the peak magnitude is a good summary of the energy in the peak. However, for broad peaks, as in JS26, the energy of the emission is distributed over many lags, and the value at a single lag captures only part of it. For such data, the value at a single lag may miss a substantial part of the emission. As an alternative, we computed the area under the curve (AUC) of the participant-level GA envelope *Ē*_*p*_(*τ* ) above the noise floor, within ±*σ*_ac_ around the reference lag:

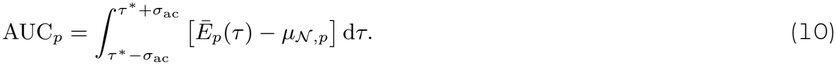

The integral was evaluated with the trapezoidal rule over lag samples. Negative values, which occur when the envelope lies below the noise floor, were set to missing. Tying the integration window to *σ*_ac_ adapts it to the expected width of the peak in each condition and makes it independent of the noisy shape of the individual peak.

The relation between AUC and peak magnitude becomes clear for a Gaussian peak of height *A* above the noise floor and width *σ*. Its full area is 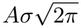. If the width of the peak matches that of the auto-correlation (*σ* ≈ *σ*_ac_), the integration window ±*σ* captures a fixed fraction 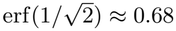 of it, so that

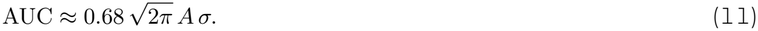

The AUC is thus proportional to the product of height and width. Conversely, 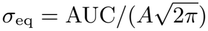 defines an equivalent width of the peak. Eq. (11) shows when the AUC adds value over the magnitude. If an experimental manipulation changes only the height of the peak, both metrics carry the same information, and the magnitude is preferable because it is simpler. If the manipulation also changes the width of the peak, or if the peak is so broad that its maximum is poorly defined, the AUC captures the change while the magnitude may miss it. Conversely, the equivalent width should be interpreted with care for noisy recordings: as *A* approaches zero, small fluctuations of the envelope within the window are divided by a small number, and *σ*_eq_ becomes inflated and unstable.

##### Peak width *σ* (pP)

Finally, we fitted the Gaussian of Eq. (8) to each participant-level GA, in the same way as described for the GA in Section 2.4.2, and used the fitted width *σ* as a metric. Unlike the previous metrics, *σ* does not depend on the overall scaling of the envelope, and it therefore describes the temporal spread of the emission independently of its strength. Since the width of the peak is dominated by the auto-correlation of the template (Section 2.4.2), changes in *σ* are expected to be small, and they are most likely to be detected for broadband templates.

##### Latency

In addition to these metrics, the cross-correlation yields the latency of the speech-like DPOAE, given by the lag of the peak. This is the key advantage of the time-domain analysis. The latency is physiologically interpretable, for instance in terms of the generation mechanisms of the emission [9, 10], and none of the frequency-domain methods in Section 2.5 provide it. Its precision, however, is limited by the width of the peak: for templates as narrowband as those of JS26, latency differences of a few milliseconds cannot be resolved.

### 2.5 Frequency-domain analysis

Cross-correlation and spectral analysis are two views of the same information. By the Wiener–Khinchin theorem, the Fourier transform of the cross-correlation between template and recording is their cross-spectrum [28]. The frequency domain therefore does not contain anything that the time domain lacks, but it offers different trade-offs. The time– frequency uncertainty principle states that a signal cannot be narrow both in time and in frequency [27]. Where cross-correlation peaks are sharp, as in MSA21 and JS25, the corresponding spectral components are broad. Where they are broad, as in JS26, the spectral components are sharply defined. Since the narrow stimulus bandwidth of JS26 forfeited much of the information in the time domain, it is natural to make use of the resulting fine-grained information in the frequency domain instead.

#### 2.5.1 DPOAE frequency band

Both spectral methods below evaluate the recording within the frequency band that the speech-like DPOAE is expected to occupy. This band can be determined in advance from the templates. For every template *w*_2*n−m*_(*t*), we computed its power spectrum with Welch’s method (Hann window of 2,048 samples, i.e. 46.4 ms, with 50 % overlap). The band was defined by the frequencies below which 5 % and 95 % of the template’s power were contained, that is, the range holding the central 90 % of its power. The band edges were averaged across all templates belonging to the same voice and stimulus condition, so that the same band was used for all trials of a condition.

#### 2.5.2 Power spectral density (PSD)

The most straightforward approach, and the one closest to the evaluation of conventional DPOAEs, is to compute the power spectral density (PSD) of the recording and to quantify the power at the expected distortion frequencies. We estimated the PSD *S*_*rr*_ (*f* ) of the recording in Pa^2^/Hz with Welch’s method [29], using Hann-windowed segments of 1 s (frequency resolution ≈ 1 Hz) with 50 % overlap. The band power of the speech-like DPOAE was obtained by integrating the PSD over the DPOAE band [*f*_l_, *f*_u_],

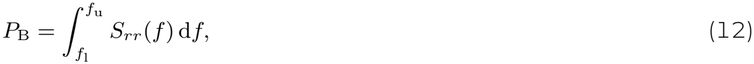

which yields a mean-square sound pressure in Pa^2^, or, expressed as a level, 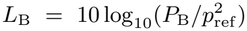 in dB SPL with *p*_ref_ = 20 μPa. The band power was computed for every trial and averaged across the trials of a participant (avgP). Applied to the three datasets, the PSD confirms the time–frequency trade-off described above: for MSA21 and JS25, the emissions are smeared over broad frequency ranges, as illustrated in Fig. 4, whereas for JS26, the four stimulus waveforms and the expected emissions appear as more sharply defined spectral peaks.

**Figure 4:**
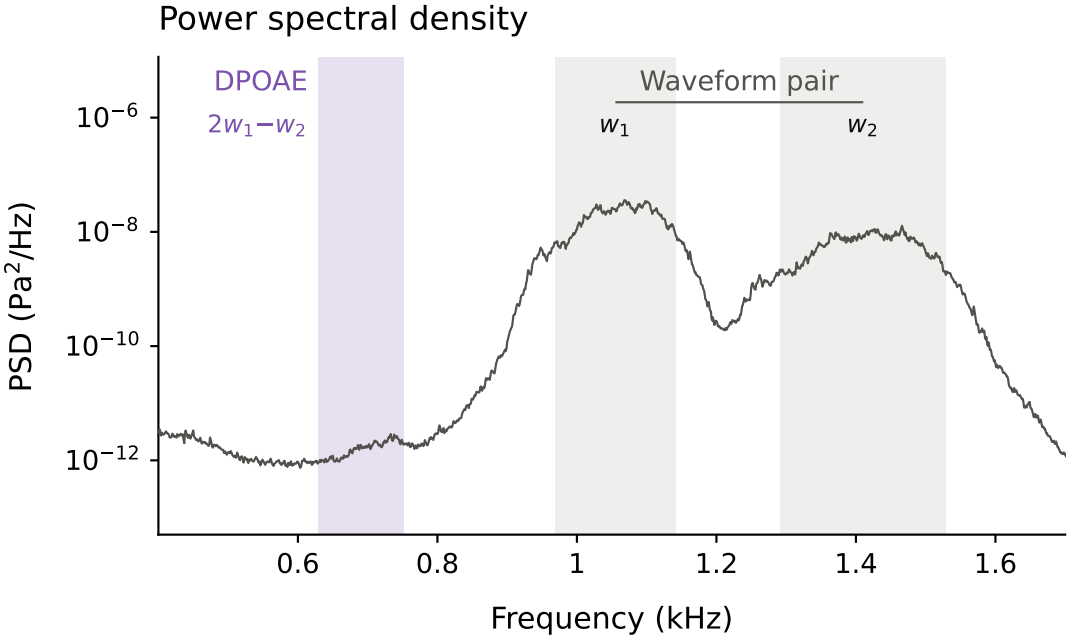
Power spectral density of an exemplary recording of the MSA21 dataset. The stimulus waveforms *w*_1_ and *w*_2_ (grey bands) follow the time-varying harmonics of the speech signal and therefore occupy entire frequency bands rather than single frequencies. The band power of the speech-like DPOAE is obtained by integrating the PSD over the band of the expected distortion product 2*w*_1_ *− w*_2_ (purple). Any noise that falls into this band contributes to the band power as well.

The PSD has, however, a fundamental limitation. It is a property of a single signal, the recording, and it does not use the template. The band power therefore contains everything that falls into the DPOAE band: the emission, but also physiological and environmental noise, and any other sound component at these frequencies. This is particularly relevant for the competing-speaker paradigm, in which the stimuli related to the competing voice may have energy in the same band. A change in band power between two conditions can thus not necessarily be attributed to a single emission unambiguously. Unlike for pure-tone DPOAEs, comparing the power at the distortion frequency with that of neighbouring frequency bins is also less straightforward, because the emission itself occupies a band rather than a single bin. We therefore include the PSD mainly as a reference for the method described next.

#### 2.5.3 Magnitude-squared coherence (MSC)

The magnitude-squared coherence (MSC) overcomes this limitation by relating the recording to the template, in the same spirit as the cross-correlation in the time domain. It is widely used in electrophysiology to quantify how strongly two signals are coupled at a given frequency [30, 31]. For two signals, here the template *w*_2*n−m*_(*t*) and the recording *r*(*t*), it is defined as

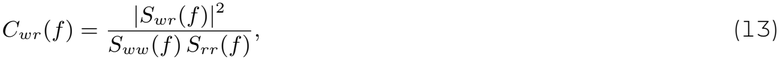

in which *S*_*ww*_ and *S*_*rr*_ denote the PSDs of template and recording and *S*_*wr*_ their cross-spectrum. The MSC lies between 0 and 1. Intuitively, it states which fraction of the power of the recording at frequency *f* can be explained by a linear transformation of the template.

This interpretation follows from a simple model of the recording. Suppose that the recording consists of the emission, which we model as the template passed through an unknown linear filter *h*(*t*) with frequency response *H*(*f* ), and of noise *n*(*t*) that is unrelated to the template:

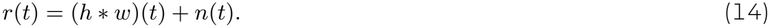

The filter captures the delay, the phase shift and the frequency-dependent gain of the emission. Then *S*_*wr*_ (*f* ) = *H*(*f* )*S*_*ww*_ (*f* ) and *S*_*rr*_ (*f* ) = |*H*(*f* )|^2^*S*_*ww*_ (*f* ) + *S*_*nn*_(*f* ), and the MSC becomes

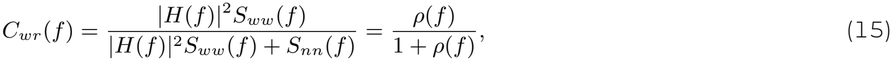

with the local signal-to-noise ratio *ρ*(*f* ) = |*H*(*f* )|^2^*S*_*ww*_ (*f* )*/S*_*nn*_(*f* ). The MSC is thus a monotonic function of the ratio between emission power and noise power at each frequency. In contrast, the PSD (Eq. 12) measures their sum, |*H*|^2^*S*_*ww*_ + *S*_*nn*_, and cannot separate the two. In particular, noise at the DPOAE frequencies increases the PSD but decreases the MSC.

Eq. (15) also shows a further property of the MSC. A time delay *τ*_d_ of the emission introduces a phase factor to the frequency response *H*(*f* ), that can be modelled as 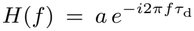 . The magnitude |*H*(*f* )|^2^ = *a*^2^ is consequently independent of *τ*_d_, meaning that the MSC is insensitive to the latency of the emission. This is an advantage in terms of robustness: unlike the time-domain metrics, the MSC does not require a reference lag, and variations of the latency across trials or participants do not reduce it. It is at the same time a limitation, because the MSC provides no information about the latency. The insensitivity holds as long as the delay is small compared with the segment length used for estimation, which is the case here by two orders of magnitude.

##### Estimation and bias correction

We estimated the MSC with Welch’s method, using Hann-windowed segments of *L* = 16,384 samples (371.5 ms) with 50 % overlap [32]. The segment length sets a trade-off. Longer segments give a finer frequency resolution, here *f*_s_*/L* ≈ 2.7 Hz, which is needed to resolve the narrow DPOAE bands of JS26. Shorter segments give more segments *K* to average over, which reduces the variance and the bias of the estimate. With the chosen value, a two-minute trial yields *K* ≈ 650 segments.

The bias arises because the MSC estimated from a finite number of segments is positive even for unrelated signals. For two independent signals, its expected value is approximately 1*/K* [32]. We therefore used a bias-corrected MSC,

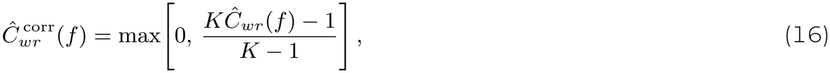

which maps the expected value for unrelated signals to zero and leaves perfect coherence unchanged. With *K* ≈ 650, the correction is small (1*/K* ≈ 0.0015), but it becomes relevant for short recordings or short segments. Since overlapping segments are not fully independent, 1*/K* slightly underestimates the actual bias.

##### Band metrics

To obtain a single value per trial, the bias-corrected MSC was averaged across the frequency bins *f*_*k*_ within the DPOAE band ℬ (Section 2.5.1). A plain band mean would treat all bins equally. However, the power of the template is not distributed uniformly across the band (Fig. 5B). At frequencies where the template has little power, there is little emission to detect, and the MSC there is dominated by noise. We therefore used a power-weighted mean,

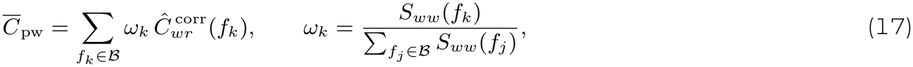

**Figure 5:**
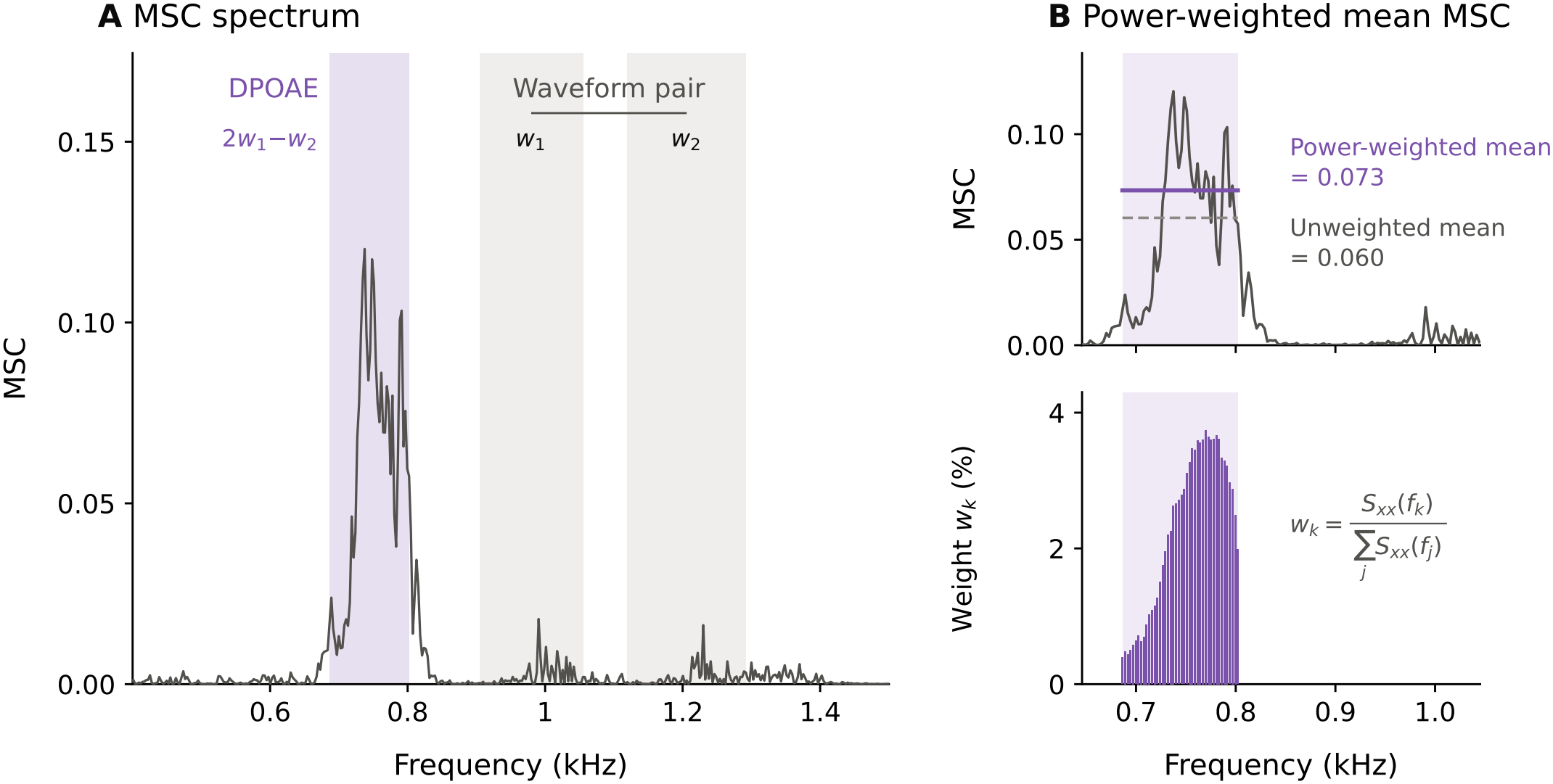
Magnitude-squared coherence (MSC) between template and recording, illustrated on an exemplary recording of the MSA21 dataset. (A) MSC spectrum. Coherence is highest within the band of the expected speech-like DPOAE (2*w*_1_ *−w*_2_, purple) and considerably lower elsewhere, including at the frequencies of the stimulus waveforms (grey). (B) Band metrics. Top: MSC within the DPOAE band with its unweighted mean (dashed) and its power-weighted mean (solid). Bottom: the weights *ω*_*k*_ are proportional to the PSD of the template, so that frequencies at which the template carries more power contribute more strongly to the power-weighted mean.

which gives more weight to frequencies that the template, and therefore presumably the emission, occupies more strongly. 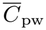 was computed for every trial and averaged across the trials of a participant (avgP). It is the MSC metric used throughout the remainder of this work, and we refer to it simply as the MSC. It can be regarded as the frequency-domain counterpart of the peak magnitude of the cross-correlation. The plain band mean and the corresponding metrics computed from the uncorrected MSC yielded very similar results and are therefore not reported.

A practical advantage of the MSC is that it does not depend on whether a peak is clearly visible. In the cross-correlation, the region in which we look for the emission is defined by the GA peak, which has to be identified first. For the MSC, the relevant frequency band is known from the templates before any data are analysed, and it can be evaluated whether or not the emission is strong.

### 2.6 Attentional modulation coefficient

For completeness, we briefly describe a derived quantity that is useful when the aim of a study is to detect attentional effects. Saiz-Alía et al. [9] quantified the effect of attention by the attentional modulation coefficient (AMC),

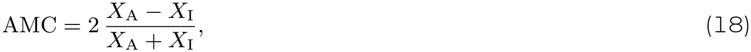

in which *X*_A_ and *X*_I_ denote the value of a DPOAE metric when the corresponding voice was attended and when it was ignored, computed per participant. The AMC is the difference between the two states relative to their mean. Positive values indicate larger values when the voice is attended, negative values larger values when it is ignored.

By setting the attended and the ignored state in direct relation to each other, the AMC removes factors that scale both states equally, such as the overall strength of a participant’s emissions, the fit of the probe, or the absolute level of the noise floor. Effects of attention can therefore become more stable across participants than when the raw values are compared. The AMC can be computed from almost any of the metrics introduced above, as long as the metric takes positive values: the peak magnitude, the AUC, the band power or the MSC. For the SNR, its use is debatable. The SNR is already scaled to the noise floor, and on the decibel scale it can be close to zero or negative, in which case Eq. (18) becomes unstable.

In the present work, we do not report AMCs. Interpreting attentional effects across all metrics and datasets is beyond the scope of this paper, and it would carry the risk of favouring metrics that happen to show strong attentional effects. Instead, we first assess how reliably each metric captures the speech-like DPOAE at all (Section 2.7).

### 2.7 Quantifying metric performance

#### 2.7.1 Separability index

##### Rationale

The metrics introduced in Sections 2.4 and 2.5 are on very different scales: band power in Pa^2^, SNR in dB, a coherence between 0 and 1, and areas and widths of the cross-correlation peak. Their raw values can therefore not be compared. Instead, we quantified the performance of a metric by how well it separates recordings that contain a speech-like DPOAE from recordings that do not. For this purpose, we compared the in-ear recordings of the participants with the out-of-ear control recordings of the same dataset (Section 2.3), which went through the identical processing pipeline. We deliberately did not define a detection threshold for the metrics, because such a threshold would be arbitrary and would depend on the units of the metric.

As a measure of separation, we used the area under the receiver operating characteristic curve (ROC-AUC), estimated non-parametrically from the Mann–Whitney *U* statistic [33–35]. We refer to it as the separability index (SI):

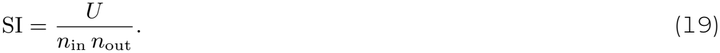

Here, *U* is the Mann–Whitney statistic of the in-ear sample, and *n*_in_ and *n*_out_ are the numbers of in-ear and out-of-ear values. The SI is the probability that a randomly drawn in-ear value is higher than a randomly drawn out-of-ear value, with ties counted as one half.

We chose this measure for four reasons: 1) It needs no detection threshold; 2) It depends only on the ordering of the values, so it is unaffected by the units of a metric and by any monotonic transformation such as linear vs. logarithmic scaling; 3) It makes no assumption about the distributions; and 4) It is directly interpretable: SI = 1 means perfect separation, SI = 0.5 means chance level, and SI *<* 0.5 means that the metric is systematically higher for the out-of-ear recordings than for the ear, i.e. that it behaves opposite to its intended direction.

It is important to keep in mind what the SI does and does not measure. A high SI shows that a metric reliably takes larger values for a recording obtained in the ear canal than outside of it. It does not show that this difference is necessarily caused by the speech-like DPOAE, nor that the metric is sensitive to physiological changes of the emission, such as those potentially caused by attention. We return to this point in the Discussion.

##### Computation

Within each dataset, trials were averaged per stimulus type per participant, without distinguishing between attentional conditions, so that each participant contributed one value per metric and stimulus type. The out-of-ear recordings were treated the same way. Missing values were removed before computing the SI. The accompanying two-sided Mann–Whitney *p*-value was stored for each metric and stimulus type.

##### Metrics and stimulus types

Six metrics entered the comparison, as defined in Section 2.4.3. Four of them were computed per trial, then averaged per participant (avgP): (1) Band power from the PSD in Pa^2^ (PSD); (2) Power-weighted, bias-corrected mean MSC (MSC); (3) Peak magnitude of the cross-correlation envelope (Mag); and (4) Signal-to-noise ratio of the cross-correlation (SNR). Two further metrics were computed on the participant-level GA (pP): (5) Area under the cross-correlation peak (AUC), and (6) Width *σ* of the Gaussian fit to the cross-correlation peak (*σ*).

SIs were computed for 14 stimulus types pooled across the three datasets: (1) MSA21 with two stimulus types, namely female and male voice; (2) JS25 with four stimulus types, namely resolved and unresolved harmonics, each with female and male voice; and (3) JS26 with eight stimulus types, namely low and transitional frequency regime, each with resolved and unresolved harmonics and with female and male voice.

The resulting 14 × 6 matrix of SIs for different stimulus types and metrics is shown as a heatmap in Fig. 6. Its diverging colour scale is centred on chance (SI = 0.5).

**Figure 6:**
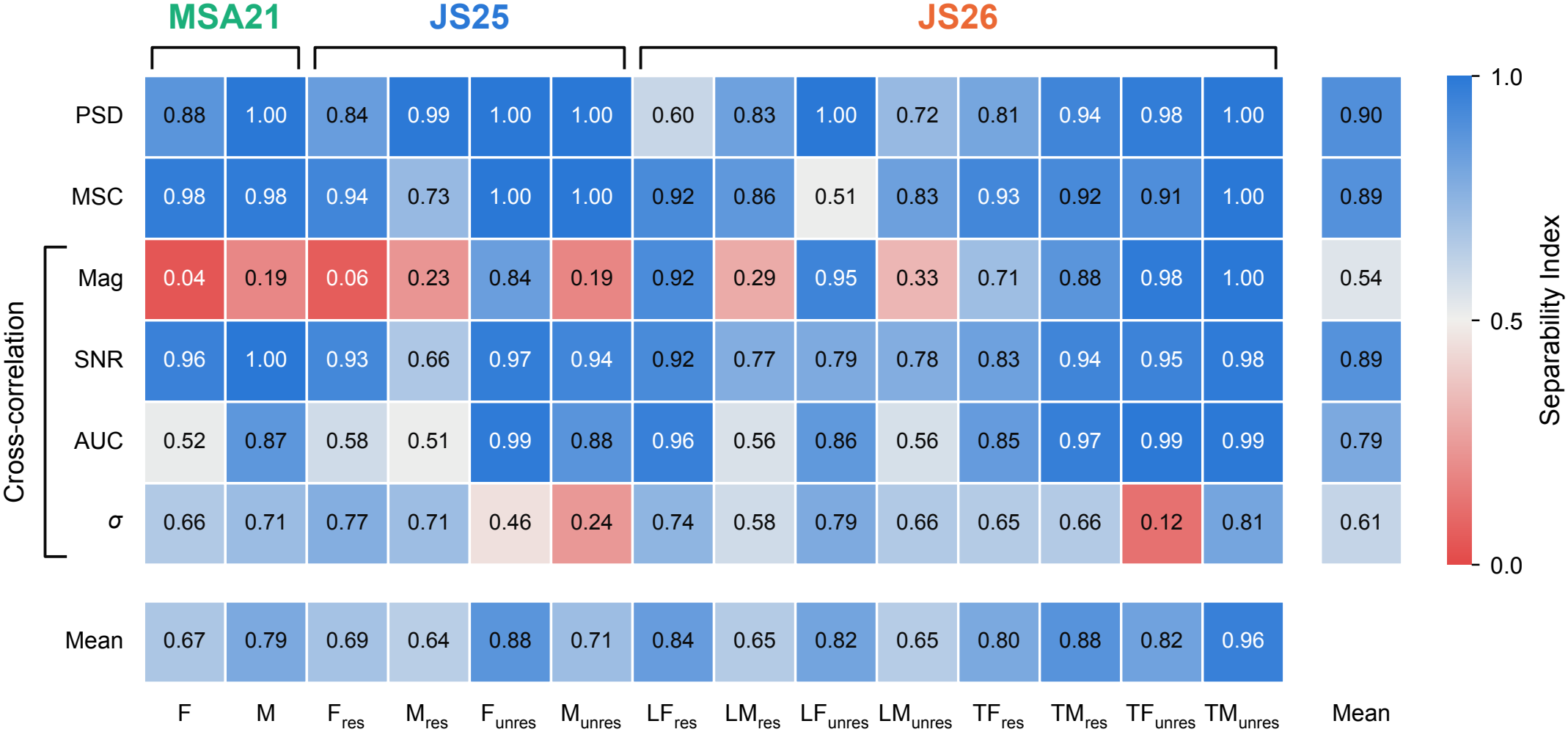
Separability index (SI) of the six metrics (rows) for the 14 stimulus types of the three datasets (columns). An SI of 1 indicates that the metric is larger for every in-ear than for every out-of-ear recording, an SI of 0.5 chance level, and an SI below 0.5 that the metric is systematically larger for the out-of-ear recordings. The bottom row and the right column give the mean SI per stimulus type and per metric, respectively. PSD: band power; MSC: power-weighted, bias-corrected magnitude-squared coherence; Mag: peak magnitude, SNR: signal-to-noise ratio, AUC: area under the curve and *σ*: peak width, all derived from the cross-correlation. Stimulus labels as in Fig. 2.

#### 2.7.2 Participant-level separability

The SI of Eq. (19) was applied to a whole group of participants. To obtain a separability value for every single participant, we applied Eq. (19) to each participant individually, that is, with *n*_in_ = 1. This participant-level SI is the fraction of out-ofear values that the participant’s value exceeds, with ties counted as one half (also known as the placement value [36]). Its mean across participants equals the group SI exactly. Participant-level SIs were computed for every participant, metric and stimulus type. They were used in two ways. To compare the metrics, they were averaged across the stimulus types of a participant, so that every participant contributed one value per metric. To compare the stimulus types, they were averaged across the six metrics, which yields one mean participant-level SI per participant and stimulus type.

#### 2.7.3 Statistical comparison of metrics

##### Omnibus test

To test whether the metrics differ in separability, we applied the Friedman test [37], with the metrics as treatments. We did so twice: with the 14 stimulus types as blocks (stimulus level, *N* = 14), and with the participants of all three datasets as blocks (participant level, *N* = 106). Within each block, metrics were ranked by SI, with average ranks assigned to ties. As the effect size, we report Kendall’s coefficient of concordance [38],

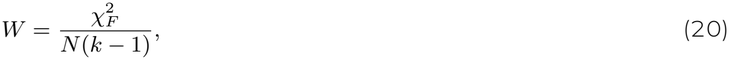

with *N* blocks and *k* = 6 metrics. *W* ranges from 0 (no consistent ranking of the metrics across blocks) to 1 (identical ranking in every block). Unlike the Friedman *p*-value, it does not depend on the number of blocks.

##### Pairwise post-hoc tests

Following a significant omnibus test, all 15 pairs of metrics were compared with two-sided Wilcoxon signed-rank tests [39] on the paired SIs. At the stimulus level, *p*-values were obtained from the exact permutation distribution of the test statistic, i.e. by enumerating all 2^*N*^ sign assignments of the paired differences, because the normal approximation is inaccurate for *N* = 14 and handles zero and tied differences poorly. At the participant level, where this enumeration is infeasible, the tie-corrected normal approximation was used. To control the family-wise error rate across the 15 comparisons, *p*-values were adjusted with the Holm–Bonferroni procedure [40] at *α* = 0.05. The results are summarised by a compact letter display [41]: metrics that share a letter do not differ significantly. Note that with *N* = 14 and 15 comparisons, the smallest attainable adjusted *p*-value at the stimulus level is about .002.

##### Critical difference diagram

To summarise the relative performance of all metrics in a single graph, we used a critical difference (CD) diagram based on the Friedman–Nemenyi procedure [42, 43], which is common practice for comparing several methods across multiple datasets. Within each block, the metrics were ranked from 1 (highest SI) to *k* = 6 (lowest SI), and these ranks were averaged across blocks to give each metric’s mean rank 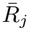. Under the null hypothesis that all metrics perform equally, every mean rank is expected to be (*k* + 1)*/*2 = 3.5. Two metrics are considered to differ significantly if their mean ranks differ by at least the critical difference

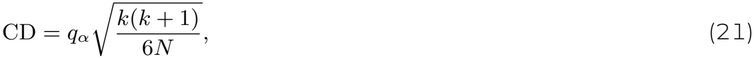

where *q*_*α*_ is the upper *α* quantile of the studentised range distribution for *k* groups and infinite degrees of freedom, divided by 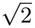. For *k* = 6 and *α* = 0.05, *q*_*α*_ = 2.850, which gives CD = 2.02 at the stimulus level (*N* = 14) and CD = 0.73 at the participant level (*N* = 106). The Nemenyi test controls the family-wise error rate by construction. In the diagram, the mean ranks are plotted on a horizontal axis, and thick bars connect groups of metrics whose mean ranks differ by less than the CD, i.e. that do not differ significantly.

The two post-hoc approaches complement each other. The Wilcoxon tests use the size of the paired SI differences and are generally more powerful. The Nemenyi test uses only the mean ranks; it is more conservative, but it gives a compact overview of which metrics form a statistically indistinguishable top group. Because the Nemenyi comparisons use mean ranks computed across all included metrics, their outcome can depend on which other metrics are part of the comparison [44]. We therefore interpret the CD diagram as a summary of the relative ranking and base conclusions about specific pairs on the Wilcoxon tests.

#### 2.7.4 Statistical comparison of stimulus types

To compare the stimulus types, we used the participant-level SIs averaged across the six metrics. All 91 pairs of stimulus types were compared. Pairs within a dataset were compared with paired Wilcoxon signed-rank tests, since they were measured in the same participants, and pairs from different datasets with Mann–Whitney *U* tests. All *p*-values were adjusted with the Holm–Bonferroni procedure and summarised by a compact letter display. In addition, Friedman tests and CD diagrams were computed within JS25 (4 stimulus types, *N* = 38) and within JS26 (8 stimulus types, *N* = 44), where every participant contributed to every stimulus type.

#### 2.7.5 Limitations of the statistical approach

The Friedman and Nemenyi tests assume independent blocks. The 14 stimulus types, however, are not fully independent: the stimulus types of a dataset were measured in the same participants and compared against the same out-of-ear recordings. Similarly, all participant-level SIs of a dataset are computed against the same 12 out-of-ear recordings. Had a different set of control recordings been obtained, the participant-level SIs of that dataset would all have shifted together. The participant-level tests treat the participants as independent observations and therefore capture the variability between participants, but not this additional uncertainty that results from the limited number of control recordings. The reported *p*-values should therefore be considered somewhat liberal.

## 3 RESULTS

### 3.1 Separability of all metrics and stimulus types

Fig. 6 shows the SI of each of the six metrics for each of the 14 stimulus types. Averaged across the stimulus types, the PSD (mean SI of 0.90), the MSC (0.89) and the SNR (0.89) separated in-ear from out-of-ear recordings best, followed by the AUC (0.79). The width *σ* (0.61) and the peak magnitude (0.54) performed considerably worse.

The metrics differed not only in their mean SI but also in how consistent they were across stimulus types. The MSC reached an SI of at least 0.73 for 13 of the 14 stimulus types; the only exception was LF_unres_ of JS26 (0.51). The SNR was even more uniform, with SIs between 0.66 (M_res_ of JS25) and 1.00, and it was the only metric that did not fall below 0.65 for any stimulus type. The AUC reached SIs of 0.84 or more for nine stimulus types, but stayed close to chance (0.51 to 0.58) for five: the female voice of MSA21, F_res_ and M_res_ of JS25, and LM_res_ and LM_unres_ of JS26.

The peak magnitude showed the largest variation of all metrics. It separated well for six stimulus types (SI of 0.84 or more: F_unres_ of JS25 and LF_res_, LF_unres_, TM_res_, TF_unres_ and TM_unres_ of JS26), but it was inverted, with an SI below 0.5, for seven of the 14 stimulus types. The inversion was most pronounced for the female voice of MSA21 (0.04) and for F_res_ of JS25 (0.06). For these stimuli, the out-of-ear recordings yielded systematically larger peak magnitudes than the in-ear recordings, even though the SNR, which is based on the same peak value, separated them well (0.96 and 0.93). The width *σ* yielded SI values between 0.46 and 0.81 for most stimulus types, but had values below 0.5 for M_unres_ of JS25 (0.24) and TF_unres_ of JS26 (0.12).

The PSD separated the in-ear from the out-of-ear recordings for all stimulus types, with SIs between 0.60 (LF_res_ of JS26) and 1.00, and reached SIs of 0.98 or more for half of the stimulus types.

### 3.2 Comparison of the metrics

The metrics differed significantly in their separability, both at the stimulus level (Friedman *χ*^2^(5, *N* = 14) = 24.9, *p <* .001, Kendall’s *W* = 0.36; Fig. 7A) and at the participant level (*χ*^2^(5, *N* = 106) = 275.7, *p <* .001, *W* = 0.52; Fig. 7B), with moderate consistency in ranking.

**Figure 7:**
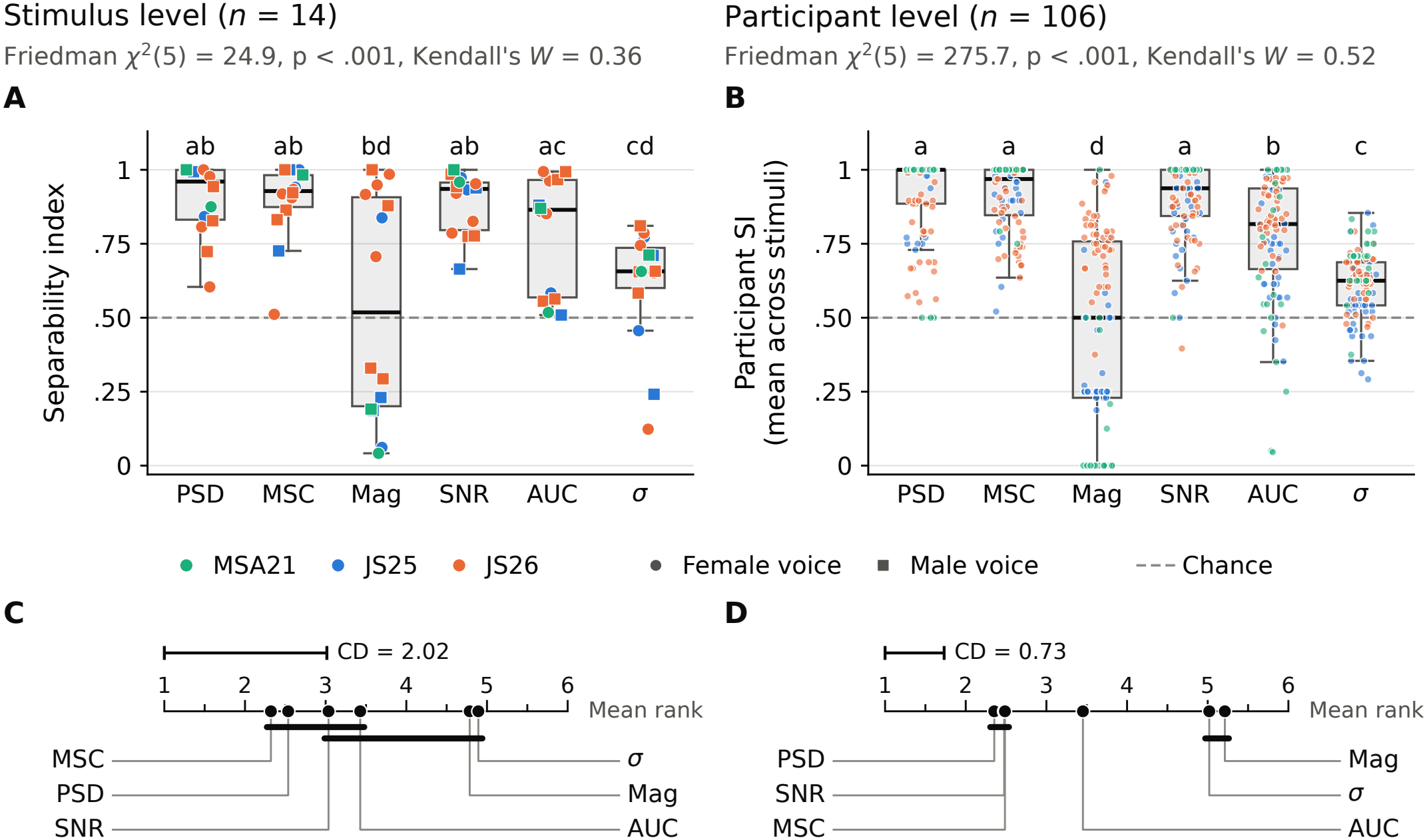
Comparison of the metrics. (A) SI per metric at the stimulus level; each point is one of the 14 stimulus types, coloured by dataset (circles: female voice, squares: male voice). (B) Participant-level SI per metric, averaged across the stimulus types of a participant; each point is one of the 106 participants. In A and B, metrics that share a letter do not differ significantly (pairwise Wilcoxon signed-rank tests, Holm-corrected). The dashed line marks chance level. (C, D) Critical difference diagrams of the mean ranks of the metrics (rank 1 = highest separability) at the stimulus level (C) and at the participant level (D). Metrics connected by a thick bar do not differ significantly (Nemenyi test, *α* = 0.05). The scale bar indicates the critical difference (CD).

At the stimulus level, the MSC obtained the best mean rank (2.32), followed by the PSD (2.54), the SNR (3.04) and the AUC (3.43), whereas the peak magnitude (4.79) and the width *σ* (4.89) ranked last (Fig. 7C). With only 14 stimulus types, few pairwise differences reached significance after Holm correction: the PSD, the MSC and the SNR separated better than *σ* (*p*_Holm_ = .009, .035 and .007), and the AUC better than the peak magnitude (*p*_Holm_ = .022). In the CD diagram, the PSD and the MSC differed from the peak magnitude and from *σ*, whereas the SNR and the AUC could not be distinguished from either the best or the worst metrics.

At the participant level, the picture became clearer. The PSD (2.35), the SNR (2.48) and the MSC (2.49) obtained the best mean ranks and did not differ from each other, but all three separated significantly better than the remaining metrics (all *p*_Holm_ *<* .001). They were followed by the AUC (3.45), which in turn separated better than *σ* (5.02) and the peak magnitude (5.21). The peak magnitude separated significantly worse than all other metrics, including *σ* (Fig. 7B and D). The participant-level distributions also show the variability within each metric: for the MSC and the SNR, most participants reached SIs above 0.75, whereas the participant-level SIs of the peak magnitude were spread over the whole range from 0 to 1, with a median close to chance.

### 3.3 Comparison of the stimulus types

The stimulus types differed markedly in how well they could be separated from the out-of-ear recordings across all metrics (Fig. 8A). Of the 91 pairs of stimulus types, 57 differed significantly after Holm correction. The highest mean participant-level SI was obtained for TM_unres_ of JS26 (0.96), which separated significantly better than every other stimulus type. It was followed by TM_res_ of JS26 (0.88), F_unres_ of JS25 (0.87) and LF_res_ of JS26 (0.84). The lowest values were obtained for M_res_ of JS25 (0.64), LM_res_ and LM_unres_ of JS26 (0.65 each), and the female voice of MSA21 (0.67).

**Figure 8:**
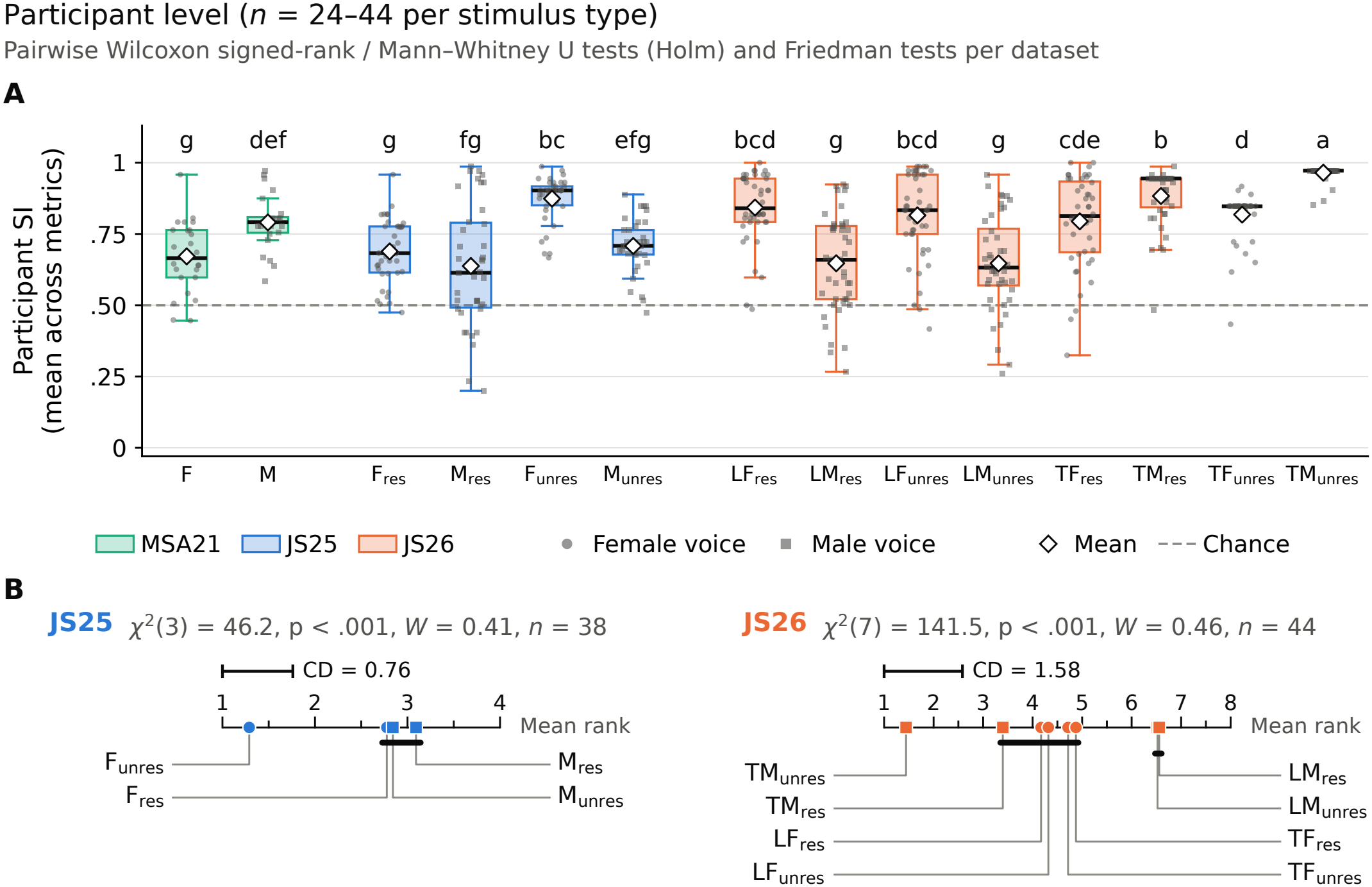
Comparison of the stimulus types. (A) Participant-level SI per stimulus type, averaged across the six metrics; each point is one participant (circles: female voice, squares: male voice), diamonds mark the mean. Stimulus types that share a letter do not differ significantly (Wilcoxon signed-rank tests within and Mann–Whitney *U* tests across datasets, Holm-corrected). The dashed line marks chance level. (B) Critical difference diagrams of the stimulus types within JS25 (left) and JS26 (right); stimulus types connected by a thick bar do not differ significantly (Nemenyi test, *α* = 0.05).

Within JS25, the stimulus types differed significantly (Friedman *χ*^2^(3, *N* = 38) = 46.2, *p <* .001, *W* = 0.41, *N* = 38; Fig. 8B). F_unres_ was clearly the best stimulus type (mean rank 1.29) and separated better than each of the other three (*p*_Holm_ *<* .001), which did not differ from each other. Within JS26, the differences were even more pronounced (*χ*^2^(7, *N* = 44) = 141.5, *p <* .001, *W* = 0.46, *N* = 44). TM_unres_ obtained the best mean rank (1.44), and the two low-frequency stimulus types of the male voice, LM_res_ and LM_unres_, the worst (6.56 and 6.52). The remaining five stimulus types formed an intermediate group. Interestingly, the voice that worked better differed between the two frequency regimes: in the low-frequency regime, the stimuli derived from the female voice separated better than those derived from the male voice, whereas in the transitional regime, the stimuli of the male voice separated better, hinting at a strong frequency dependence of the emission strength.

## 4 DISCUSSION

We compared six metrics for the evaluation of speech-like DPOAEs, four in the time domain and two in the frequency domain, across three datasets with 14 stimulus types and 106 participants. As a common criterion, we used how reliably each metric separates in-ear recordings from out-of-ear control recordings. Overall, the PSD, the MSC and the SNR separated in-ear from out-of-ear recordings most reliably, followed by the AUC, whereas the peak magnitude and the peak width *σ* were the least reliable. For the PSD, however, this high separability mainly shows that the measurement setup worked as intended, not that the emission was captured, as discussed below. At the same time, the performance of every metric depended on the stimulus type, and the stimulus types themselves differed considerably in how well their emissions could be measured. There is therefore no single metric that suits every experiment. In the following, we discuss what the separability index tells us, what the strengths and weaknesses of the individual metrics are, and how the results can guide the design and analysis of future experiments.

### 4.1 Interpretability of the separability index

The SI quantifies how reliably a metric takes larger values in the ear than outside of it. This is a necessary condition for a metric that is meant to capture an otoacoustic emission, but not a sufficient one: a metric can potentially separate in-ear from out-of-ear recordings on other grounds than the DPOAE. Conversely, a low SI does not necessarily imply that a metric fails to capture the emission. The SI of a metric should therefore always be interpreted together with the properties of the recordings it is applied to.

The PSD illustrates the first point most clearly. It reached the highest mean SI of all metrics, although the band power contains everything that falls into the DPOAE band (Section 2.5.2). In a sealed ear canal, the sound pressure produced by the probe is much higher than in the free field, and with it the spectral leakage of the stimulus waveforms, physiological noise and any other sound component in the DPOAE band. A larger band power in the ear is therefore expected even in the complete absence of an emission, and, consistent with this, the separability of the PSD hardly depended on the frequency of the emission (Section 4.6). Level-based metrics such as the band power are, moreover, directly affected by any difference in the settings of the measurement chain between in-ear and out-of-ear recordings, whereas metrics that relate a quantity to a reference from the same recording, such as the SNR and the MSC, are not. A high band power should therefore not be taken as a marker for the presence of speech-like DPOAEs, but rather as a sanity check, confirming that the probe sat in the ear canal and that the stimuli were delivered and recorded as intended.

The peak magnitude shows that both cases can occur for the same metric. For LF_unres_ of JS26, the grand average did not show a clear peak (GA SNR of 2.0 dB; Fig. 2) and the MSC was at chance level (SI = 0.51), so that no reliable emission, or at least no strong one, probably occurred. Nevertheless, the peak magnitude (0.95) and the PSD (1.00) separated in-ear from out-of-ear recordings almost perfectly. These high SIs therefore reflect differences that are unrelated to the emission: the influence of noise in the case of the PSD, and the normalisation of the cross-correlation in the case of the peak magnitude (Section 4.2). Conversely, the peak magnitude separated poorly for MSA21, although its grand averages showed clear peaks with the highest GA SNRs of all datasets, and it reached an SI of only 0.06 for F_res_ of JS25, although the participant data showed a robust GA peak and the out-of-ear recordings none (Fig. 2).

Although the peak magnitude, unlike the band power, is based on the template of the DPOAE, a comparison with out-of-ear recordings is prone to error for both metrics, and their SIs say little about whether an emission was captured unless the grand average is taken into account. No metric should therefore be trusted blindly. The SI is best used alongside the grand average and an understanding of what the metric represents in a given design.

### 4.2 Peak magnitude

The peak magnitude, the metric used in previous work [10], yielded SIs that varied most across stimulus types and were even below 0.5 for half of them, including F_res_ of JS25, for which the participant data showed a robust GA peak (GA SNR of 6.6 dB) and the out-of-ear recordings did not show any. For these stimuli, the out-of-ear recordings yielded larger magnitudes than the in-ear recordings, although the SNR, which is based on the same peak value, separated them well. In the out-of-ear recordings, the whole envelope of the cross-correlation was therefore elevated, not only its value at the reference lag.

We attribute this mainly to the free-field condition of the out-of-ear recordings, which affects the values of the normalised cross-correlation (Eq. 4). The normalisation divides by the energy of the entire recording. In a sealed ear canal, the recording is dominated by the stimulus waveforms, which are much louder than the emission and than the noise in the DPOAE band. Chance correlations between the template and the noise in its band are therefore divided by a large number and result in small correlation coefficients. With the probe in the free field, the sound pressure of the stimulus waveforms at the microphone is much lower, so that noise in the DPOAE band makes up a larger share of the recording, and the same chance correlations result in larger coefficients. Where the emission is strong, as for F_unres_ of JS25 or the transitional stimuli of JS26, the in-ear peak still exceeds this offset. Where it is weak, as for the low-frequency stimuli of the male voice, the offset dominates. The SNR is not affected, because the peak and the noise floor are divided by the same factor.

For MSA21, the SI value below 0.5 is more surprising, since the DPOAEs for these stimuli were strong, with the highest GA SNRs of all datasets (18 to 19 dB). We cannot determine its cause with certainty, but the most likely explanation is that the in-ear and the out-of-ear recordings of MSA21 were obtained with different recording chains. The participant data were collected in London, with stimulus presentation and recording controlled by MATLAB, whereas the out-ofear recordings were obtained years later in Erlangen, with the setup and software used for JS25 and JS26. Even with matched levels, differences in sound card settings, preamplifier noise or filtering change what a recording consists of, and thereby the normalisation of the cross-correlation. MSA21 is the only dataset in which in-ear and out-of-ear recordings differ in this respect. The stimulus level, in contrast, is unlikely to be the cause: all three datasets used the same average stimulus level of 57 dB SPL, and the peak magnitude nevertheless separated well for several stimulus types of JS25 and JS26 (SI of 0.84 or more for F_unres_ of JS25 and LF_res_, TM_res_, TF_unres_ and TM_unres_ of JS26). Moreover, the GA magnitudes of MSA21 were no larger than those of JS25 (Fig. 2), although its GA SNRs were considerably higher. An emission that is strong relative to the noise is not necessarily large relative to the whole recording, and this is precisely the difference between the SNR and the peak magnitude. A change in the recording chain affects neither the SNR nor the MSC, since both relate the emission to a reference from the same recording. The case of MSA21 therefore underlines that the peak magnitude alone is not suited to distinguish in-ear from out-of-ear recordings, and that the grand averages always have to be examined first, in particular when parts of the recording chain may have changed between recordings.

A low SI of the peak magnitude therefore does not mean that the metric is unsuited to DPOAE detection. Rather, the comparison with free-field recordings is not a fair test for a metric that depends on the total energy of the recording, and certain conditions need to be verified before the metric can be trusted. The most important of these concerns the grand average: the participant data should show a robust GA peak, and the out-of-ear recordings should not show any. For F_res_ of JS25, both conditions were met, so that the peak magnitude can be expected to capture the emission despite its low SI. Conversely, a high SI of the peak magnitude, as for LF_unres_ of JS26, does not indicate an emission if the GA peak is absent.

The dependence on the total energy of the recording has a further practical consequence. The peak magnitude does not only depend on the emission, but on everything that changes the energy of the recording, such as the fit of the probe, the level of the stimuli in the ear canal, or the volume of the ear canal. This is one reason why the magnitudes vary so much between participants and stimulus types, as is also apparent from the grand averages (Fig. 2). The compared conditions should therefore be recorded with the same setup and the same acoustic input, so that these factors are identical, and the GA peak should be stable across all compared conditions. Both requirements were met in JS25 and JS26: the attended state of one voice was recorded at the same time as the ignored state of the other voice, the acoustic input was identical in all attentional conditions, and the analysis was restricted to stimulus types with clear GA peaks [10]. For comparisons between stimulus types, participants or datasets, however, the peak magnitude is not suited.

### 4.3 Signal-to-noise ratio

The SNR relates the peak magnitude to the noise floor of the same trial and thereby compensates for the scaling factors discussed above. Accordingly, it was among the best metrics and one of the most uniform across stimulus types. Its main limitation lies in what the noise floor represents. The noise floor does not only contain acoustic noise, but also chance correlations between the template and the stimulus waveforms, including those of the competing voice. These correlations differ between templates. A higher SNR of one stimulus type compared with another can therefore result from a lower noise floor rather than from a stronger emission. The SNR is thus well suited to establish whether an emission is present, and to compare conditions that share the same template, but it is not an effective tool to compare speech-like DPOAEs between different stimulus types.

### 4.4 Magnitude-squared coherence

Among the metrics that make use of the template, the MSC was the most reliable. It obtained the best mean rank at the stimulus level, did not differ from the SNR and the PSD at the participant level, and reached SIs of at least 0.73 for all stimulus types except LF_unres_ of JS26, for which, as discussed above, there is no evidence for a reliable emission. Several properties of the MSC contribute to this. It is evaluated directly from the recording and the DPOAE template, without a grand average, a reference lag or a fit. The frequency band in which the emission is expected is known in advance from the template, so that no peak has to be detected first. As a ratio between coherent and total power (Eq. 15), it is insensitive to factors that scale the whole recording. And it is insensitive to the latency of the emission and to variations of the latency across trials. Consistent with these properties, the separability of the MSC depended less on the frequency of the emission than that of the AUC and the SNR (Section 4.6). This makes the MSC the most suitable metric for comparing speech-like DPOAEs across stimulus types, participants and datasets. Its main limitation is the counterpart of its robustness: it provides no information about the latency of the emission.

### 4.5 Area under the curve and peak width

The AUC was designed for broad cross-correlation peaks, and it performed accordingly. For most stimulus types of JS26, whose templates were narrowband and whose peaks were broad, it reached high SIs. For the narrowband peaks of JS25, the integration window of ±*σ*_ac_ is short, and the AUC then offers little beyond the peak magnitude; for F_res_ and M_res_ of JS25, its SIs stayed close to chance. The AUC is therefore a useful complement to the peak magnitude when the peaks are broad, in particular because it is less sensitive to the exact choice of the reference lag.

The width *σ* was among the least reliable metrics. This is not surprising. As shown in Section 2.4.2, the width of the peak is dominated by the auto-correlation of the template, which is determined by the stimulus design and is therefore the same whether or not an emission is present. In addition, the fitting procedure itself favours widths close to that of the auto-correlation, since *σ*_ac_ serves as the starting value of the fit and determines the window within which it is performed. In-ear and out-of-ear recordings therefore tend to yield similar widths. Moreover, the SI assumes that larger values indicate a stronger emission, which is not necessarily true for a width. A Gaussian fitted to a peak-less, noisy envelope can yield an arbitrary, often large width, which explains the inverted SIs for M_unres_ of JS25 and TF_unres_ of JS26. We therefore regard *σ* as a descriptive quantity of the peak shape, rather than as a metric to detect an emission.

### 4.6 Implications for stimulus design

The stimulus types had at least as much influence on the resulting emissions as the metrics with which we characterized them. Stimuli derived from unresolved harmonics in the transitional regime of JS26 (TM_unres_) and from unresolved harmonics of the female voice in JS25 (F_unres_) produced the most reliable emissions, whereas stimuli whose emissions lay at low frequencies performed worst. In an exploratory analysis, the mean participant-level SI of a stimulus type, averaged across the six metrics, correlated with the mean frequency of its DPOAE template (Spearman’s *ρ* = 0.79, *p <* .001, *N* = 14). All stimulus types with emissions below about 700 Hz, namely M_res_ of JS25, LM_res_ and LM_unres_ of JS26, and the female voice of MSA21, were among the least reliable. This is in line with the frequency dependence of speech-like DPOAEs observed in JS25, where amplitudes were largest around 1 kHz and the emission at 350 Hz could hardly be measured [10], with the frequency dependence of pure-tone DPOAEs [2], and with the increase of physiological and equipment noise towards low frequencies [3]. Across the individual metrics, the dependence on emission frequency was strongest for the AUC (*ρ* = 0.83) and the SNR (*ρ* = 0.61), weaker for the MSC (*ρ* = 0.55) and the peak magnitude (*ρ* = 0.49), and weakest for the PSD (*ρ* = 0.34), consistent with the band power reflecting the recording conditions rather than the emission.

This correlation should not be over-interpreted. The 14 stimulus types are not independent, and the frequency of the emission is confounded with several other properties that differ between the datasets and that we cannot disentangle in retrospect. These include the number of simultaneously presented stimulus pairs, which determines the correlations between stimulus waveforms and templates and thereby the noise floor; the bandwidth *B*_0_ of the fundamental waveform; the distance between the two voices, ΔERB; whether the stimuli were derived from resolved or unresolved harmonics, which in turn is linked to the frequency range in which a given harmonic number lands; and the ratio *m/n* of the stimulus pair, which, however, was similar across all datasets. In particular, the small ΔERB of JS26 may be relevant for attentional effects, which we discuss elsewhere [11], but it is not reflected in the separability of the emissions.

The bandwidth *B*_0_ deserves a separate comment, since it is a design choice with a clear trade-off. A narrow band-width, as in JS26, reduces the correlations between the stimulus waveforms and the templates, but it forfeits temporal resolution: the cross-correlation peaks become broad, and the latency of the emission can no longer be determined precisely. A wide bandwidth, as in MSA21 and JS25, preserves temporal resolution and with it the latency information, at the cost of broader spectral bands. The choice of *B*_0_ should therefore be guided by whether the latency is of interest. If it is, broadband templates and a time-domain analysis are needed. If it is not, narrowband templates combined with the MSC are a robust alternative.

If the goal is simply to measure speech-like DPOAEs as reliably as possible, the present results suggest designing stimuli similar to TM_unres_ of JS26 or F_unres_ of JS25, with emissions between roughly 1 and 2.5 kHz. In most studies, however, the stimulus design is constrained by the research question, for instance by the requirement to target resolved harmonics of a voice. In such cases, the stimulus type in Fig. 6 that is most similar to the intended design can serve as a guide to which metric is likely to evaluate the data most reliably.

### 4.7 Comparison with existing methods

The analyses of speech-like DPOAEs published so far relied on the peak magnitude of the cross-correlation. Saiz-Alía et al. [9] quantified attentional effects by the AMC computed from the peak magnitude, and Steinebach and Reichen-bach [10] compared the peak magnitudes of attended and ignored states directly. Our results show that the peak magnitude is the least robust of the metrics considered here. This does not invalidate the comparisons made in these studies. They compared conditions that shared the same template and that were recorded with the same setup and the same acoustic input, so that factors that scale the whole recording affected all compared conditions alike. This also holds for F_res_ of JS25, the stimulus type with the strongest attentional effects: its low SI results from the free-field out-of-ear recordings, whereas the GA showed a robust peak in the participant data and none in the out-of-ear recordings. The AMC used in MSA21 additionally relates the attended to the ignored state, which removes such scaling factors to a large extent (Section 2.6). Nevertheless, the low separability of the peak magnitude for several stimulus types means that any conclusion based on it requires a stable GA peak in all compared conditions. The SNR is the more robust alternative and preferable for future analyses. The MSC is the corresponding alternative in the frequency domain. It requires no prerequisites, such as a clear GA peak, to be meaningful and, in addition, allows comparisons across stimulus types.

## 5 CONCLUSIONS

Speech-like DPOAEs can be evaluated both in the time and in the frequency domain, and the best choice of metric depends on the stimuli and on the question to be answered. Across three datasets, the magnitude-squared coherence and the signal-to-noise ratio of the cross-correlation separated in-ear from out-of-ear recordings most reliably among the metrics that make use of the expected waveform of the emission. The band power separated equally well, but since it contains every contribution in the DPOAE band regardless of its origin, it can only serve as a sanity check of the measurement setup, not as a marker for the presence of an emission. The MSC is particularly suited to compare speech-like DPOAEs across stimulus types, since it can be evaluated directly from the recording without prerequisites, whereas the cross-correlation remains indispensable when the latency of the emission is of interest. The peak magnitude, used in previous work, depends on the total energy of the recording and should only be used for comparisons within the same recording session and with a stable grand average peak. Stimuli whose emissions lie between roughly 1 and 2.5 kHz yielded the most reliable measurements. We hope that the methods and guidelines presented here, together with the openly available code, make speech-like DPOAEs more accessible and encourage their use in further studies of cochlear processing of natural sounds.

## DATA AVAILABILITY

All three datasets analysed in this study are publicly available:

- MSA21 [9], on figshare: https://doi.org/10.6084/m9.figshare.12738515
- JS25 [10], on Zenodo: https://doi.org/10.5281/zenodo.16837673
- JS26 [11], on Zenodo: https://doi.org/10.5281/zenodo.24104371

## CODE AVAILABILITY

All custom code used for stimulus generation, analysis of speech-like DPOAEs and statistical evaluation is openly available on GitHub at https://github.com/janna-stb/speechlike_dpoae_followup under the MIT License.

## AUTHOR CONTRIBUTIONS

J.S.: Conceptualisation, Methodology, Software, Investigation, Formal analysis, Visualisation, Writing – original draft; T.R.: Conceptualisation, Supervision, Resources, Funding acquisition, Writing – review & editing.

## DECLARATION OF GENERATIVE AI AND AI-ASSISTED TECHNOLOGIES IN THE MANUSCRIPT PREPARATION PROCESS

During the preparation of this work, the authors used Claude (Anthropic) to formulate, revise and edit the text of the manuscript. In addition, Claude was used as a supporting tool during the development of the analysis code, for instance to revise, extend and debug existing code. All code was reviewed, tested and validated by the authors. After using this tool, the authors reviewed and edited the content as needed and take full responsibility for the content of the published article.

